# Model-based tumour subclonal deconvolution accounting for spatio-temporal sampling biases

**DOI:** 10.1101/2025.03.31.646463

**Authors:** Elena Rivaroli, Elena Buscaroli, Salvatore Milite, Alberto Casagrande, Giulio Caravagna

## Abstract

Bulk DNA sequencing has entered the clinic, and understanding tumour evolution in space and time has become critical for advancing precision oncology. A recent work by us has brought a population genetics perspective into the classical tumour subclonal deconvolution problem, showing how multiple spatiotemporal sampling biases arise when we collect more than one sample of the same tumour. Sampling biases complicate the mapping of the mutations into the evolutionary process, a complex issue undermined by existing multi-sample analysis methods. This work presents `MOBSTERm`, a novel mixture-within-mixture Bayesian framework that extends our earlier approach to a multi-dimensional formulation. Our model incorporates distinct mathematical distributions that capture sampling bias patterns in tumour data, allowing the deconvolution to resist the effect of some confounders. This works presents the simulation of one source of sampling bias analysed with this new approach, together with a large scale test based on simulations. Moreover, we apply our model to understand subclonal deconvolution from a multi-region whole-genome colorectal cancer sample and from longitudinal whole-genome glioblastoma samples of `15` patients collected before and after treatment. This new deconvolution approach offers a better account of the effect of spatio-temporal tumour biases, allowing us to better elucidate complex clonal dynamics from multi-sample cancer sequencing data.

**Code availability:** `MOBSTERm` is available at the GitHub page https://github.com/caravagnalab/MOBSTERm. The code to reproduce our analyses is available at Zenodo https://doi.org/10.5281/zenodo.14867256.

## 1 Introduction

Bulk DNA sequencing is a routine tool in clinical oncology, enabling large-scale tumour genome profiling [1]. Genetic mutations revealed by sequencing can inform of selective pressures that ultimately shape clonal architectures, drive phenotypic diversity, and inform tumour evolutionary patterns [2]. The field is slowly moving to collect multi-region and longitudinal datasets to measure tumour samples from different locations (primary/metastasis) and time points (pre-treatment/post-treatment).

In this era, the complexity of genomics data is only tamed by computational frameworks reconstructing tumour evolution across space and time [3–6]. This new generation of tools already provides significant advancement to the field, yet with some limitations for multi-sample datasets, which can be subject to several sampling biases caused by our inability to sequence whole tumours. In our recent work [7], we have proposed a model-based machine learning method to integrate tumour subclonal deconvolution with population genetics, a theoretical framework that models the interplay of mutation, selection, and drift during evolution. While this model was resolute in showing the opportunity to integrate a mathematical model with machine learning, in the same paper, we discovered that most existing methods often fail to address the spatial and temporal dimension of tumour evolution, where clonal interactions and selective pressures vary across regions. This observation stems from considering mutations conditionally independent given the latent clustering structure, a common approach of most methods [3–6]. Our work established three sources of spatial bias [7]. The hitchhiker’s mirage arises when mutations are neutral in one region and hitchhikers in another. The admixing deception occurs when a sample contains genetically distant lineages, producing multiple variant allele frequency (VAF) peaks that mimic selection, but result from lineage mixing. Finally, the ancestor effect mimics the sampling of random ancestors from a tree, which do not correspond to actual clones with distinct evolutionary parameters.

This study introduces a novel Bayesian framework that integrates model-based population genetics with a multi-dimensional formulation to overcome at least part of these challenges. By adopting a mixture-within-mixture modelling approach, our framework enables a more robust reconstruction of clonal architectures and evolutionary trajectories from multiple samples, resisting some of the confounders introduced in [7]. We validate our approach through extensive evaluations of simulated and real-world datasets, demonstrating its ability to allow a more simple interpretation of mutation data from multi-region assays compared to state-of-the-art methods. Our results highlight the power of computational methods to decode tumour complexity, paving the way for more effective precision oncology strategies using multi-sample assays.

## 2 Multivariate subclonal deconvolution

Following our earlier work on `MOBSTER`, we introduce a Bayesian multivariate extension of the approach that combines population genetics and machine learning to perform bulk DNA deconvolution; we refer to this model as `MOBSTERm`.

### A mixture-within-mixture problem

Subclonal deconvolution methods employ mixture models to identify distinct subpopulations within bulk cancer samples, where each cluster correspond to a clone. Traditional univariate mixture models for tumour subclonal deconvolution assume the independence of the *n* input mutations [3–7]. Their canonical multi-dimensional extension can process mutations across *d* ≥1 dimensions (i.e., multiple samples), while still assuming the independence across the dimensions [3–6]. Given *k* mixture components, the general likelihood of such models is

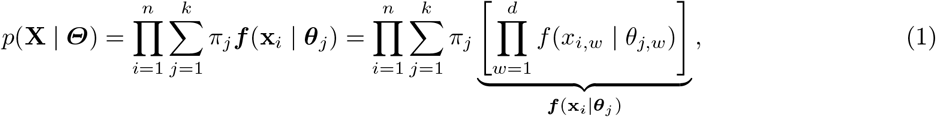

where **X** = {**x**_1_,…, **x**_*n*_} represents the set of mutations, and each mutation **x**_*i*_ = ⟨*x*_*i*,1_,…, *x*_*i,d*_⟩ is a *d*-dimensional measurement. Each cluster is parametrised by a vector ***θ***_*j*_ = ⟨*θ*_*j*,1_,…, *θ*_*j,d*_⟩, whereas the mixing proportions ***π*** are a canonical *k*-dimensional stochastic vector such that *π*_*j*_ ≥ 0 and 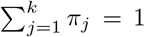 This broad family of models includes several tools [3–6], where ***f*** typically follows a Binomial or Beta-Binomial likelihood to model the variant read counts of somatic mutations (Fig. 1a). In these models, the clustering responsibilities, i.e., the posterior distributions that define the probability of each mutation belonging to a cluster, are defined by (*n* × *k*)-dimensional latent variables

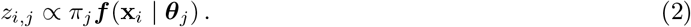

**Fig. 1:**
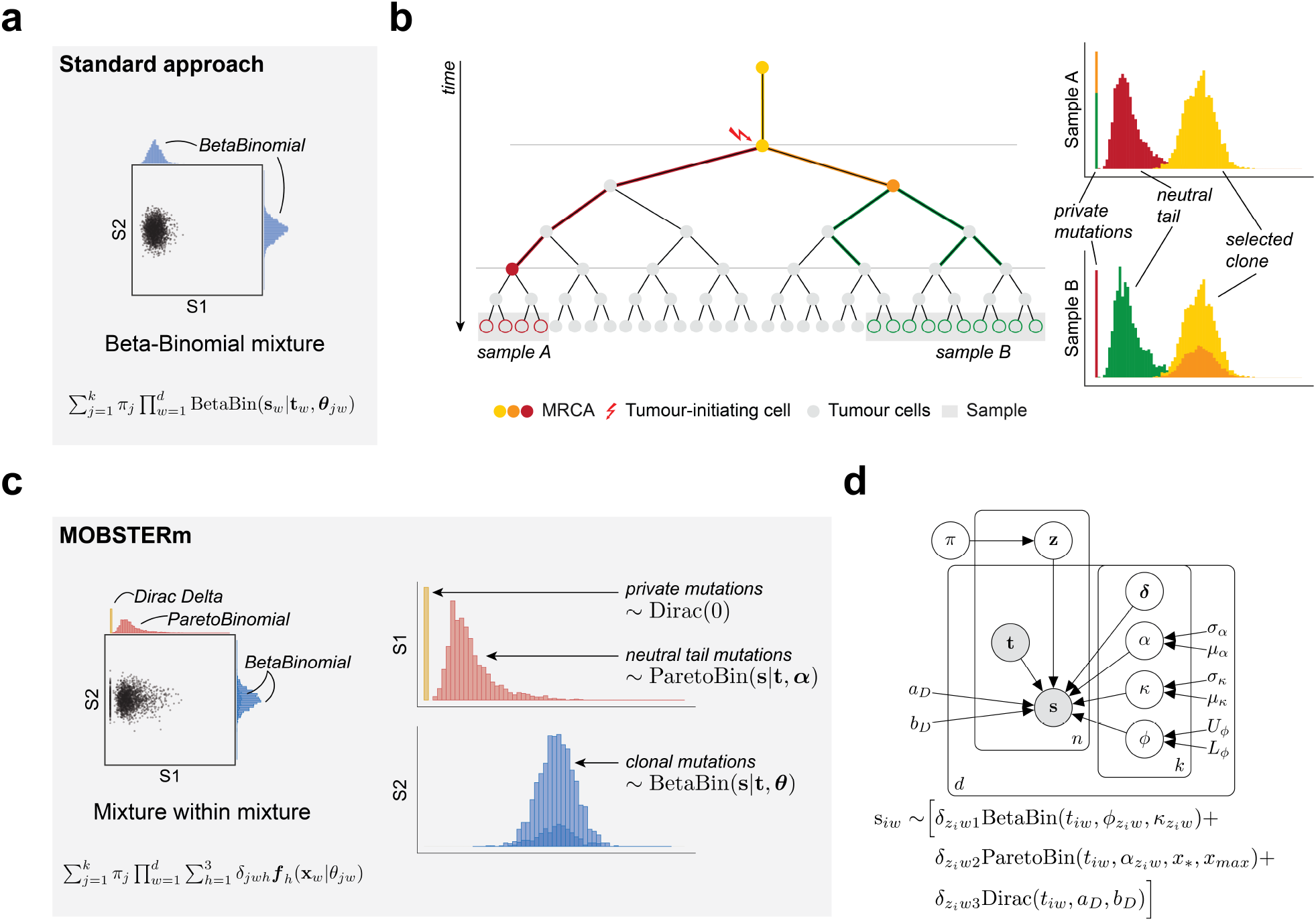
**a**. Standard multivariate subclonal deconvolution models a mixture of equivalent distributions (e.g., Beta or Beta-Binomial), with independent parameters across dimensions. **b**. Tumour phylogenetic tree with a neutral expansion, where two biopsies (sample A and B) are collected. Mutations in the yellow branch are clonal, i.e., shared between samples; in the orange branch, they are clonal in B (absent in A); in the green branch, they are neutral for sample B and absent in sample A; in the red branch, are neutral for A and absent in B. In multivariate subclonal deconvolution, these mutations must be modelled using three different distributions: a power-law for neutral (tail) mutations, a peaked non-zero for clonal (or subclonal) mutations, and a Dirac Delta for private mutations. The same mutation can follow distinct distributions in A and B. **c**. `MOBSTERm` achieves this with a mixture-within-mixture model. For each dimension of each component, we model read counts as a mixture of three possible distributions (Beta-Binomial, Pareto-Binomial and Dirac), allowing the model to identify which distribution best fits the data. **d**. Probabilistic graphical model representing observed (grey nodes) and latent variables for `MOBSTERm`. The observed number of reads with the variant **s** is distributed as a mixture of Beta-Binomial, Pareto-Binomial and Dirac distributions, weighted through a parameter ***δ***.

In our recent work [7], we have augmented this class of models by introducing `MOBSTER`, a frequency-based maximum likelihood approach that uses a mixed version for *f*. The inspiration of `MOBSTER` is from population genetics which predicts that the frequency *f* of passengers mutations in an expanding population follows a Landau distribution [8], whose asymptotic behavior can be approximated by a power-law distribution 1*/f* ^2^ [8–15]. Besides introducing this hybrid mixture, in [7], we have observed how spatial sampling biases can confound the inferences of standard models. In particular, one key confounder that we identified, termed hitchhiker’s mirage, shows that in some cases, the marginal data distribution (in multiple dimensions) might follow distinct likelihood types (which can be known). This happens when mutations are neutral in one sample but hitchhiking in another, which can be exacerbated when we collect multiple samples in time or space.

We, therefore, extend the model in equation (1), allowing every dimension to follow a distinct likelihood function, with the caveat that we cannot know, a priori, the likelihood type per dimension. We achieve this by introducing *k* new latent variables, one per dimension, as a tensor in *k* × *d* × 3 dimensions and adopt a mixture-within-mixture approach

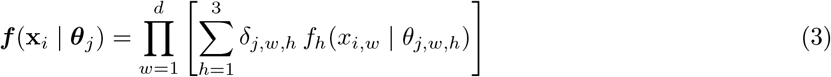

where ***δ*** is the *k* × *d* × 3 matrix of mixing proportions for the dimension-specific distribution assignments, such that *δ*_*j,w,h*_ ≥ 0 and 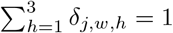

In this model, we extend the frequency-based approach of `MOBSTER` [7], with three possible density functions *f*_*h*_: the Beta-Binomial, Pareto-Binomial and Dirac Delta (Fig. 1b).

These densities model, respectively, hitchhikers, neutral mutations and zero-frequency mutations, i.e., mutations private to one or more samples (Fig. 1c). The mathematical definitions of these densities are introduced in the following paragraphs.

### Density distributions

Let the input data **X** consist of *n* pairs of values, each describing independent experiments, that is **X** = [**s, t**]. Here, **s ∈** ℕ^*n*×*d*^ represents the number of variant reads covering a given somatic mutation, while **t** ∈ ℕ^*n*×*d*^ denotes the sequencing depth at the corresponding locus. In this model, the number of variant reads (**s**) and the sequencing depth (**t**) are modelled as the number of successes and trials in a Binomial distribution, respectively. The Beta-Binomial is a standard distribution to model read counts from mutations hitchhiking in subclones under positive selection [3–5], which uses a Beta prior over the Binomial probability of success to model sequencing overdispersion. Using the *ϕ* ∈ [0, 1] and *κ >* 0 parametrization for the Beta distribution, the shape parameters *a >* 0 and *b >* 0 are expressed as *a* = *ϕ κ* and *b* = (1 − *ϕ*)*κ*. Given ***ϕ*** and *κ* and total number of trials **t** *>* 0, the Beta-Binomial density for mutation *i* and dimension *w* is

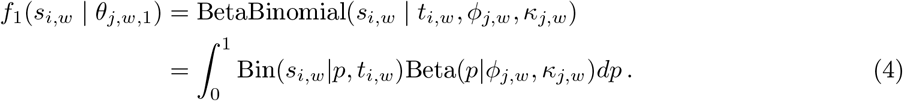

A novel distribution specifically introduced in `MOBSTERm` is the Pareto-Binomial, a compound distribution in which a Pareto prior is placed on the probability of success in the Binomial distribution. As the Pareto distribution models the variant allele frequencies (VAFs) of mutations coming from a neutral tail, the Pareto-Binomial distribution models read counts of neutral mutations. In addition to the scale parameter *x*_*_, representing the VAF lower bound, and the shape parameter ***α*** *>* 0, we also augment the Pareto with a right truncation at *x*_max_ = *n*_*B*_ *ρ/*(*n*_*A*_ + *n*_*B*_), where *ρ* is the sample purity, and *n*_*A*_ and *n*_*B*_ represent the major and minor allele copy numbers, respectively. In the current version of `MOBSTERm`, it is assumed that all cells within a sample share the same copy number state, meaning that *n*_*A*_ and *n*_*B*_ are fixed for each sample. The right truncation avoids frequencies above the clonal peak [7]. Given *x*_*_, ***α*** *>* 0 and *x*_max_, the Pareto-Binomial density is

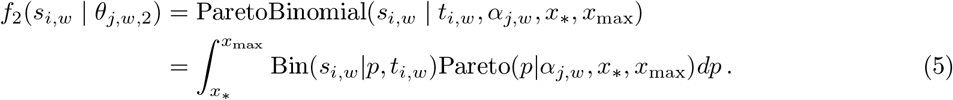

The current implementation of `MOBSTERm` contains some mathematical simplifications. First, we set a detectability threshold on allelic frequencies at 3%, leading to *x*_*_ = 0.03. Second, to avoid the complexity of computing the integral, we approximate the Pareto-Binomial distribution by sampling a probability value *p* from a Pareto distribution and then using this value as the probability of success in a Binomial distribution.

Finally, the Dirac distribution models zero-frequency mutations, with a mass concentrated at a single point, hereby *x* = 0, to model mutations with 0 mutant reads. Its theoretical density is

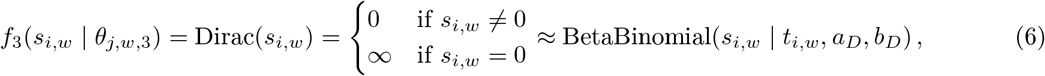

which, in practice, we approximated by a mutant distribution sharply peaked around *x* = 0 to avoid computing infinite values. We fixed the parameters for this distribution to *a*_*D*_ = 10^−4^ and *b*_*D*_ = 10^4^. This corresponds to a Beta with mean 10^−8^ and variance 10^−12^. Given this mean and variance, even when used as a prior in a Binomial distribution, the probability of observing a non-zero value remains extremely low.

### Complete Bayesian model

Given the input data **X**, the model’s parameters ***θ*** and the mixing weights ***π*** and ***δ***, the likelihood of our model is

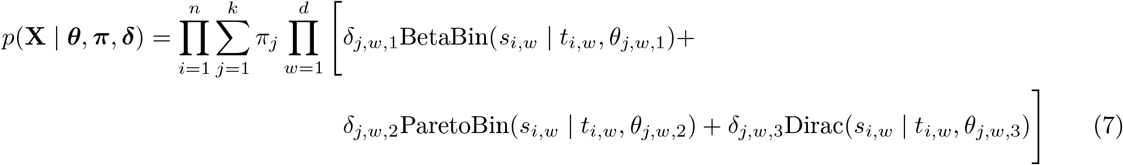

where ***θ*** = (***ϕ, κ, α***, *x*_*_, *x*_*max*_, *a*_*D*_, *b*_*D*_), with *x*_*_, *a*_*D*_, *b*_*D*_ and *x*_max_ being fixed parameters. The `MOBSTERm` joint distribution (Fig. 1d) is factored as

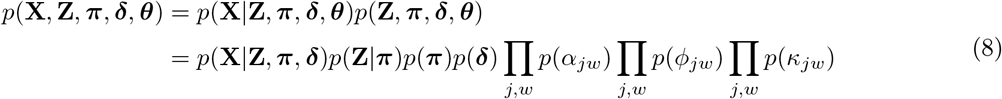

where the prior distributions are

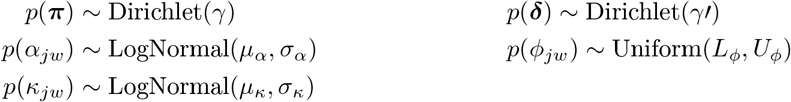

and the fixed-value hyperparameters are *µ*_*α*_, *σ*_*α*_, *L*_*ϕ*_, *U*_*ϕ*_, *µ*_*κ*_, and *σ*_*κ*_. In our analyses, we set *µ*_*α*_ = log(1.5) and *σ*_*α*_ = 0.5, based on prior biological knowledge of the Pareto distribution shape parameter [8–15]. Similarly, we set *L*_*ϕ*_ = max(*x*_*_, 0.1) and *U*_*ϕ*_ = *n*_*A*_ *ρ/*(*n*_*A*_ + *n*_*B*_), reflecting the theoretical VAF peaks for clonal and subclonal clusters. Finally, we set *µ*_*κ*_ = log(200), and *σ*_*κ*_ = 0.01, based on empirical observations of the Binomial variance.

### Model implementation

The parameters of `MOBSTERm` are learnt using Stochastic Variational Inference (SVI), implemented in the probabilistic programming language Pyro [16]. In this framework, the objective is to minimize the evidence lower bound (ELBO)

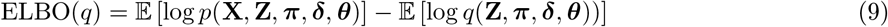

with a variational distribution *q*(·). Once the parameters are learnt, we select the optimal number of clusters *k* through model selection, employing an extension of the Bayesian Information Criteria (BIC) termed the Integrated Completed Likelihood (ICL) [17]. The ICL is defined as

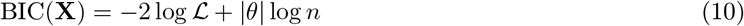

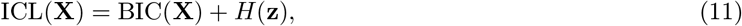

where ℒ is the likelihood computed at the Maximum A Posteriori (MAP) estimates of the parameters, and *H*(**z**) is the entropy of the clustering assignments **z**

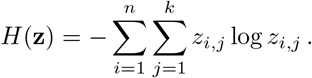

## 3. Results

### 3.1 Experimental evaluation of the performance of **MOBSTERm**

We sampled datasets from the generative model, varying the number of mutations, true clones and samples. Specifically, we generated datasets with 5000, 10000 and 15000 mutations, and 2, 3 and 4 samples. In each generated dataset, we sampled one clonal and one neutral tail cluster from a Beta-Binomial and a Pareto-Binomial, respectively, with the same distribution across all samples. Additional clusters were randomly sampled to create datasets with 4, 6, and 8 total clusters. In these clusters, observations for each dimension were generated independently following one of three possible distributions, i.e., Beta-Binomial, Pareto-Binomial, or a value of 0 to represent private mutations. The sampled mixture weights determine the number of mutations assigned to each cluster. For each combination of parameters, we generated 15 datasets. We simulated data with purities of 70, 90 and 100% and sequencing depth of 70x and 100x, and compared the ability of our model to assign mutations to clusters against PyClone-VI [3], a tool that in our earlier work [7] showed good performance (Fig. 2).

**Fig. 2:**
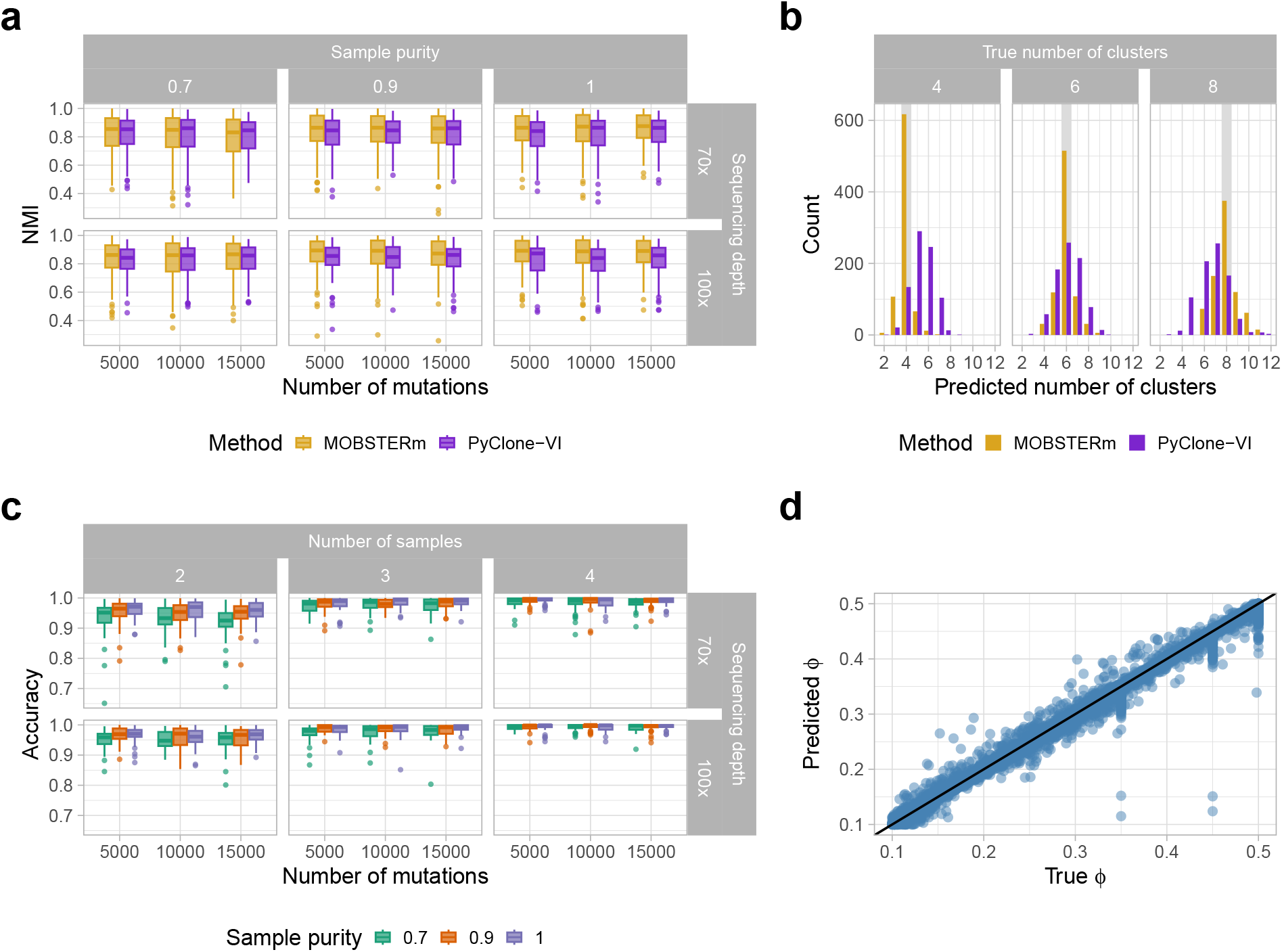
Results on simulated data. **a**. Normalized Mutual Information (NMI) comparison between `MOB-STERm` (orange) and PyClone-VI (violet) [3] for synthetic datasets. The results show the reached performance increasing number of mutations, sample purity and sequencing depth, aggregating datasets with varying number of samples and cluster. **b**. Comparison of the true vs. predicted number of clusters *K* across all the generated datasets. Each plot corresponds to a different true number of clusters (*K* = {4, 6, 8}). The histogram shows the predicted *K* values distribution. These results follow from single-sample analysis in [7], where models that control for neutral tails identify fewer clusters. **c**. Accuracy of predicted distribution types (Pareto-Binomial, Beta-Binomial, or Dirac) assigned to individual mutations compared to their true distribution, increasing number of mutations, sample purity and sequencing depth, and aggregating datasets with varying number of samples and cluster increasing sample purity and sequencing depth. **d**. True and predicted values of the centroid parameter *ϕ*, for the components following a Beta-Binomial distribution across all synthetic datasets.

We measured clustering performance with Normalized Mutual Information (NMI) (Fig. 2a). Across all tests, we observed strong performance with median NMI values greater than 0.8, suggesting our model’s ability to assign mutations to clusters accurately. As expected, the performance improves with the number of mutations, as well as with the sample purity and coverage, proving how increasing the signal for each component and reducing its noise increases the ability of the model to detect the clusters correctly. Additionally, the NMI decreases as the number of clusters increases. This is likely due to more overlapping components, leading to noisier observations and to increased complexity in cluster identification. In most of the tested configurations, PyClone-VI showed a lower NMI and inferred a different number of clusters than the true number (Fig. 2a,b). Since PyClone-VI does not directly model Pareto-distributed tails, the fits are confounded by neutral mutations. Thus, similar to what is observed in [7], the method often fits extra Beta-Binomial clusters to fit neutral tails. Performance metrics stratified for number of samples and number of clusters are shown in Supplementary Figures 1-6.

We further evaluated `MOBSTERm` ability to correctly identify the distribution type of each mutation (Pareto-Binomial, Beta-Binomial, or Dirac). `MOBSTERm` showed good performance in reconstructing the components’ distributions, consistently achieving overall median accuracy values above 0.9. As expected, accuracy improved with increased dataset sizes, dimensionality, sample purity and sequencing depth (Fig. 2c). Notably, `MOBSTERm` showed higher accuracy in identifying Beta-Binomial and Dirac distributions compared to Pareto-Binomial ones (Supplementary Fig. 7). Nevertheless, even in this cases, the median accuracy remained greater than 0.7 across all tests.

Finally, we evaluated the estimated values of the Beta-Binomial clusters centroid *ϕ*, compared to the true ones (Fig. 2d). The predicted and actual values show a strong correlation, showing that our model is not only able to retrieve mutation clusters and distributions correctly, but it can recover the correct centroids.

### 3.2 Identifying the hitchhiker’s mirage with **MOBSTERm**

In [7], we introduced the “hitchhiker’s mirage” as a cause for the overestimation of subclones in multi-sample analyses. This confounder describes neutral mutations spreading across different tumour regions, mistakenly classified as selected subclonal mutations by current deconvolution methods. This happens easily in polyclonal tumours with two monoclonal samples (Fig. 3a), where a founder clone accumulates neutral mutations that are low frequency in one sample, but hitchhike at a high frequency in a second sample (with the subclone). These mutations follow a Pareto tail in the first sample, and a Beta-Binomial in the second, and a standard method will identify them as a subclonal expansion, suggesting a mistaken linear evolution pattern.

**Fig. 3:**
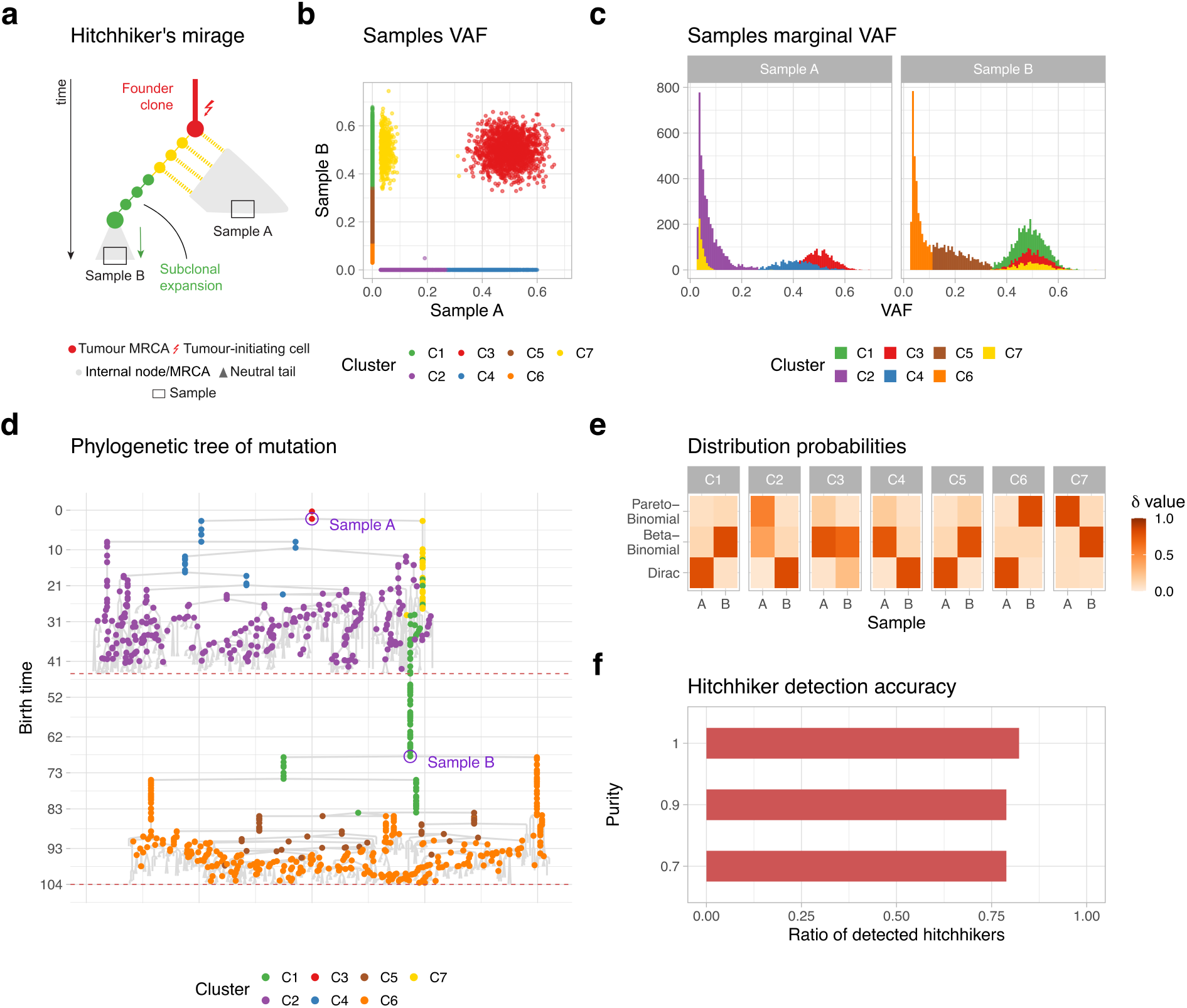
Hitchhiker’s mirage confounder. **a**. Illustration of the hitchhikers mirage confounder. **b, c**. Fit of our model on one of the simulated hitchhikers mirage. **d**. Phylogenetic forest of the mutations sequenced in the RACES [18] simulation. The mutations has been highlighted according to the `MOBSTERm` assignments, showing the true phylogenetic relationship among the identified components. **e**. Heatmap of the values of ***δ*** parameters for each sample of each component found. The hitchhiker component (cluster C7) has been correctly classified with high certainty to a Pareto-Binomial distribution in sample A and Beta-Binomial in sample B. **f**. Accuracy in detecting the hitchhiker’s mirage in 135 datasets generated with RACES [18], with different purities of 70, 90 and 100% and sequencing depth of 50x, 70x and 100x. The accuracy is computed as the ratio of correctly identified hitchhiker components across all samples. The results are aggregated by coverage values, with separate results for each coverage shown in Supplementary Fig. 8.

We sought to use `MOBSTERm` to analyse tumours where this confounder was present. We used a simulation environment for spatial tumour evolution (RACES [18], in preparation) and then fitted the simulated data with `MOBSTERm`. The model was able to correctly identify the hitchhiker cluster, i.e., cluster C7 (Fig. 3b,c). The identification of this cluster helps us to state that the observed cluster does not correspond to an additional subclone but rather arises due to the hitchhiking of neutral mutations. In real case datastets, the neutrality of hitchhiking clusters should be further supported by the absence of driver mutations. These results are further supported by Fig. 3d, which shows the tree associated with the RACES simulation, with mutations mapped on it, coloured based on the clustering assignments. The hitchhiker cluster is correctly assigned to a Pareto-Binomial distribution in sample A and to a Beta-Binomial in sample B with high certainty (Fig. 3**e**). We extended the tests by combining purities of 70, 90 and 100% and sequencing depth of 50x, 70x and 100x. In most of the simulations, the model was able to correctly identify an hitchhiker cluster, i.e., a Pareto-Binomial distribution in one sample and a Beta-Binomial distribution in the other sample (Fig. 3f).

### 3.3 Analysis of glioblastoma and colorectal adenocarcinoma cancer data

#### Longitudinal glioblastoma samples

We analysed high-resolution whole-genome sequencing (WGS) samples collected before and after treatment for 15 glioblastoma patients [19]. These data had median coverage around 100x and were previously analysed in [7]. For this analysis, we focused on regions with a single karyotype, following [7]. Our analysis revealed 5 cases with no evidence of subclonal expansion and 8 cases with a subclone, present in primary and/or relapse samples (Supplementary Fig.9), consistently with earlier analyses [7, 19]. As expected, and as observed in the results on synthetic datasets, the Dirac assignment are generally more confident, while some uncertainties remains in distinguishing between Pareto-Binomial and Beta-Binomial distributions. In two cases, mutations expected to follow a Beta-Binomial were instead fit to a long Pareto-Binomial tail (patients H043-D9MRCY and H043-GKS176). Although our model struggled detecting these additional clusters due to the low number of mutations, it helped to disentangle the true distributions with greater uncertainty in the classification between Pareto-Binomial and Beta-Binomial components. Finally, cases where a possible subclone mixed with the neutral tail were instead flagged by uncertainty in the ***δ*** assignment.

For patient H043-BU96 (Fig. 4a-d), whose samples have 9153 mutations, our analysis revealed a clonal cluster of shared mutations, neutral mutations private to each sample, as well as a shared cluster of mutations that appear neutral in both samples, classified as Pareto-Binomial in both primary and relapse. In this patient it was instructive to inspect the uncertainty between Pareto-Binomial and Beta-Binomial assignments in the private neutral tail of the primary sample, possibly caused by a small VAF peak ∼0.15 that could appear as a subclone. The analysis revealed the presence of a subclone in the relapse sample, reported also in [7], where it has been identified a driver mutation LINC00689, corroborating the hypothesis of a subclone in the relapse sample. Additionally, the model identified a subclonal peak at high frequency in the primary sample. However, since there are no identified driver mutations, this could be due to a sampling bias not handled by `MOBSTERm`. Accounting for confidence levels in the identified clusters, we also constructed a clonal tree (Fig. 4d), helping us to distinguish potential true subclones from sampling biases.

**Fig. 4:**
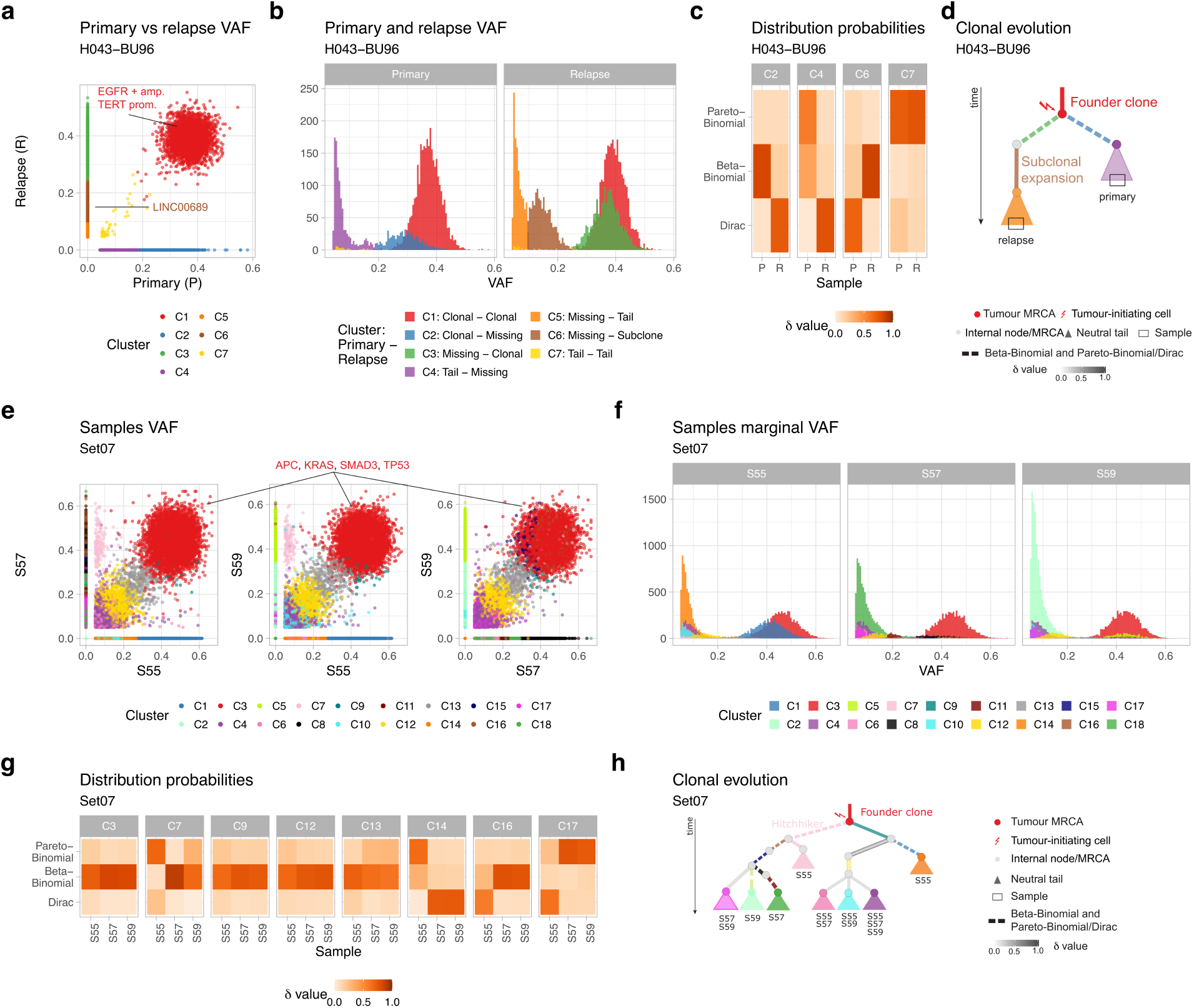
Analysis of longitudinal glioblastoma and multi-region colorectal adenocarcinoma tumours. **a**,**b**. `MOBSTERm` fit on patient H043-BU96 from the analysed glioblastoma cohort. Results identified a positively selected subclone private to the relapse sample, which contains a driver mutation in LINC00689. **c**. Heatmap of the ***δ*** parameter for selected clusters from patient H043-BU96, indicating the probabilities assigned to each possible distribution type for individual clusters in each sample. Cluster C4 shows uncertainty between Pareto-Binomial and Beta-Binomial in the primary sample. The heatmap for all clusters is shown in Supplementary Fig. 9. **d**. Clone tree for patient H043-BU96. The branch leading to the primary sample is characterized by a neutral evolution, showing a mild uncertainty about the presence of an additional subclone. The branch leading to the relapse sample presents a plausible subclonal expansion. In both samples a neutral expansion is present, which gives us the possibility to identify an MRCA for each sample. **e**,**f**. `MOBSTERm` fit on samples S55, S57 and S59 of the Set07 dataset. Note the samples present an example of the hitchhiker’s mirage, identified by the method as component C7. **g**. Heatmap of the ***δ*** parameter for selected clusters for samples S55, S57 and S59 of Set07. Here, component C7, the hitchhiker, is assigned to a Pareto-Binomial distribution in sample S55 with high certainty, and to a Beta-Binomial in the other samples. Clusters C12 and C13 have been classified to Beta-Binomial distributions but shows some uncertainty towards Pareto-Binomial ones. Cluster C9 has been classified to Beta-Binomial in all samples, however the low number of mutations assigned to it suggests that these mutations may be part of the clonal cluster C3. The heatmap for all clusters is shown in Supplementary Fig. 10. **h**. Clone tree for Set07. Cluster C7, is clonal in both S57 and S59, but is found in the neutral tail in S55, providing evidence it is not due to a subclonal expansion but rather to the hitchhiker confounder. Note how clusters C12 and C13 report uncertainty in the distribution assignment, indicating they might be indeed neutral tails.

#### Multi-region colorectal cancer samples

We applied our method to high-resolution multi-region WGS data from a colorectal cancer with 100x median coverage (Fig. 4e-h). We analysed 3 samples from the Set07 dataset, previously analysed in [7], with a purity of 0.88 and 45237 total mutations. `MOBSTERm` identified 18 total clusters, most of which contain private mutations assigned to a Dirac distribution in one or more samples. The method detected four clonal clusters of shared mutations (C3, C9, C12 and C13). Clusters C12 and C13 are composed of a small number of mutations and exhibit high uncertainty in the ***δ*** parameters, suggesting that these mutations may belong to a misclassified neutral tail, as proposed in [7, 20]. Furthermore, only clonal driver mutations APC, KRAS, SMAD3 and TP53 were identified (Fig. 4e). The absence of subclonal drivers in these clusters [7, 20] further supports the hypothesis that clusters C12 and C13 represent neutral tails. We additionally identified clonal clusters and neutral tails private to single samples and private clusters at high frequency, possibly due to spatial confounders described in [7]. Interestingly, in these samples is present a group of hitchhiking mutations, present at high frequency in samples S57 and S59, and at low frequency in sample S55. `MOBSTERm` is able to identify a cluster of hitchhiking mutations (C7). This component is classified as a Pareto-Binomial distribution in sample S55, and as a Beta-Binomial in samples S57 and S59. This result confirms in a real-case scenario how `MOBSTERm` is able to identify the hitchhiker’s mirage, reporting also a level of confidence of the assigned distributions (Fig. 4g). After fitting the model, we reconstructructed a clonal evolution tree accounting for uncertainties in potential subclones and neutral tails (encoded in the ***δ*** parameters), while highlighting the presence of the hitchhiker confounder on tumour evolution (Fig. 4h).

## 4 Conclusion

This study expands on our previous work [7] to introduce `MOBSTERm`, a Bayesian framework for interpreting spatio-temporal evolutionary trajectories from multiple cancer samples. `MOBSTERm` combines model-based population genetics with a multidimensional approach, enhancing the inference of tumour evolutionary dynamics across various spatial and temporal samples.

Recently, there has been increasing debate about interpreting subclonal deconvolution patterns from single samples [21, 22], and the focus is shifting toward aggregating signals from multiple samples. However, this collection can introduce subtle sampling biases that complicate data interpretation. This works offers a well-grounded solution to the hitchhiker’s mirage, a statistical confounder first highlighted in [7]. In this work, by means of extensive synthetic validations, we have shown that `MOBSTERm` is robust and accurate to resolve the hitchhiker’s mirage confounder. In the application with real data, we have also shown that after the fit with the model, we are able to construct a clonal evolution tree which includes uncertainty about potential subclones or neutral tails. This is the first attempt to bring population genetics in the multivariate deconvolution problem.

While `MOBSTERm` is innovative, its underlying algorithm can still be improved. The main limitation of the model is its assumption that all mutations within each sample share the same karyotype. While addressing this limitation would increase the model’s complexity, it would also enhance its ability to capture tumour heterogeneity and improve the accuracy of subclonal reconstruction. Moreover, our previous work identified two spatial confounders related to tumour ancestor sampling. These confounders may lead to an overrepresentation of clonal lineages that are not linked to distinct evolutionary parameters, complicating subclonal deconvolution. A deeper understanding of these confounders is necessary for their effective integration into mixture models, also considering mathematical techniques other than density estimation. Overall, our findings highlight the need to combine mathematical models of evolution with machine learning. As we gain better access to sequencing across different regions and over time, we need tools that can handle sampling biases and clonal changes to improve precision in cancer treatment. Our approach is a step forward, laying the groundwork for future work that can include more evolutionary factors, single-cell details, and functional data to enhance our understanding of tumour development.

## Acknowledgments

The research leading to these results has received funding from AIRC under MFAG 2020 - ID. 24913 project – P.I. Caravagna Giulio. We acknowledge financial support under the National Recovery and Resilience Plan (NRRP), Mission 4, Component 2, Investment 1.1, Call for tender No. 1409 published on 14.9.2022 by the Italian Ministry of University and Research (MUR), funded by the European Union – NextGenerationEU– CUP J53D23015060001.

## Disclosure of Interests

The authors declare no competing interests

## Notes

### Competing Interest Statement

The authors have declared no competing interest.

