## Supplementary figures for "Model-based tumour subclonal deconvolution accounting for spatio-temporal sampling biases"

### 1 Supplementary figures

Supplementary Fig. 1: Normalized Mutual Information (NMI) comparison between MOBSTERm (orange) and PyClone-VI [1] [1] (violet) for synthetic datasets with purities of 70% and sequencing depth of 70x. The performance improves with the number of mutations as well as with the number of samples, while decreases as the number of clusters increases, likely due to more overlapping components.

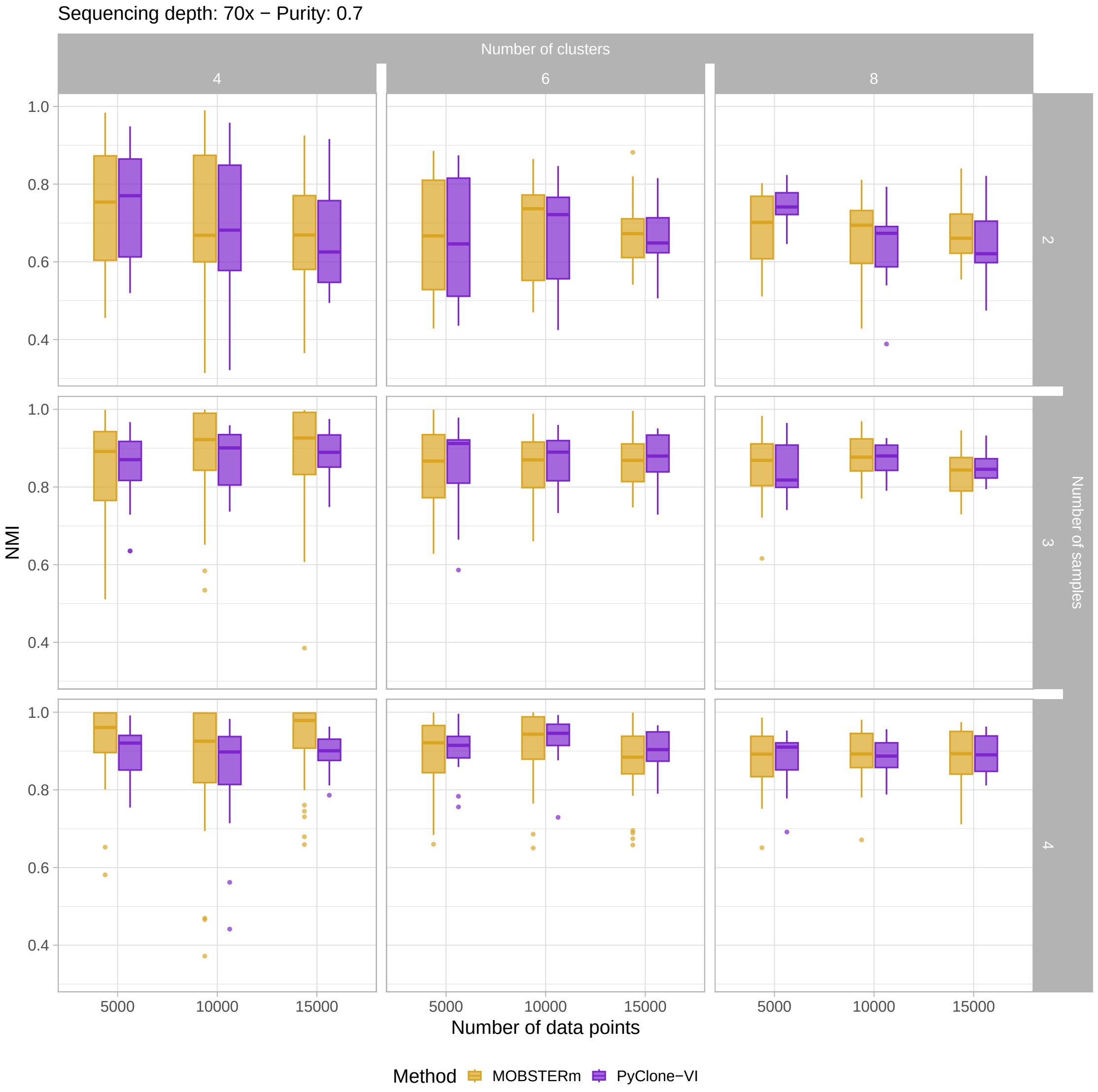

Supplementary Fig. 2: NMI comparison between MOBSTERm (orange) and PyClone-VI [1] (violet) for synthetic datasets with purities of 90% and sequencing depth of 70x.

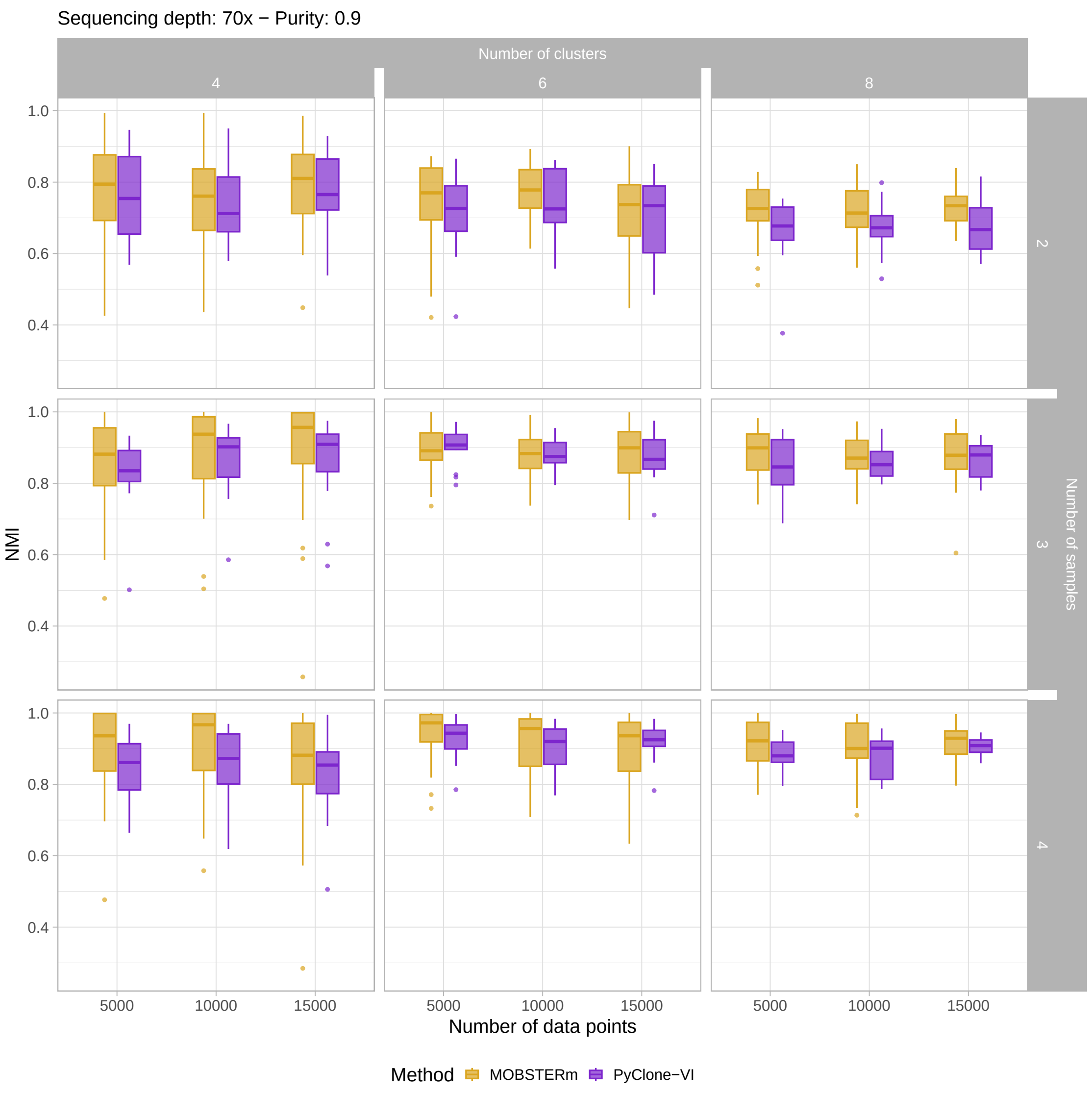

Supplementary Fig. 3: NMI comparison between MOBSTERm (orange) and PyClone-VI [1] (violet) for synthetic datasets with purities of 100% and sequencing depth of 70x.

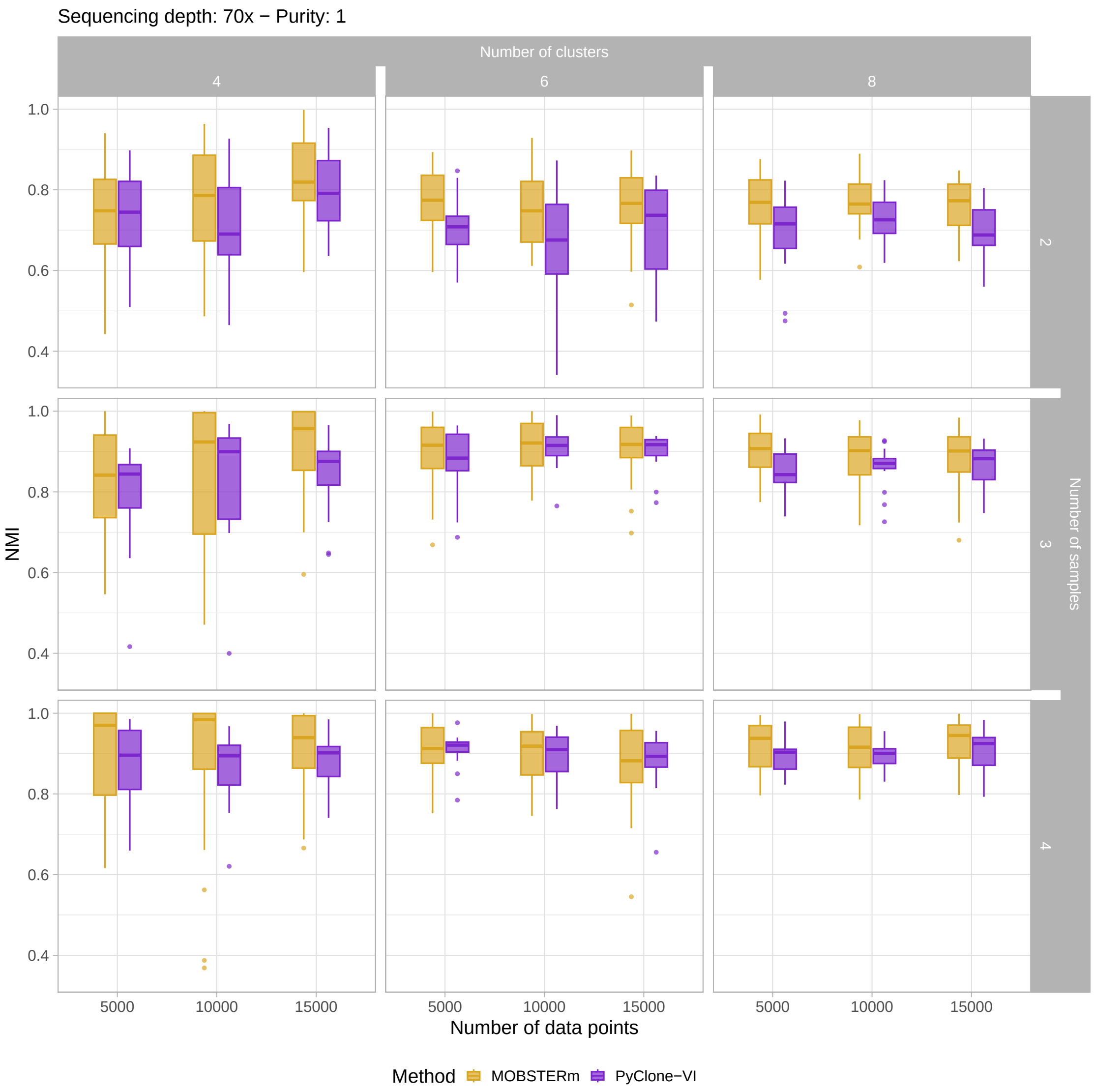

Supplementary Fig. 4: NMI comparison between MOBSTERm (orange) and PyClone-VI [1] (violet) for synthetic datasets with purities of 100% and sequencing depth of 100x.

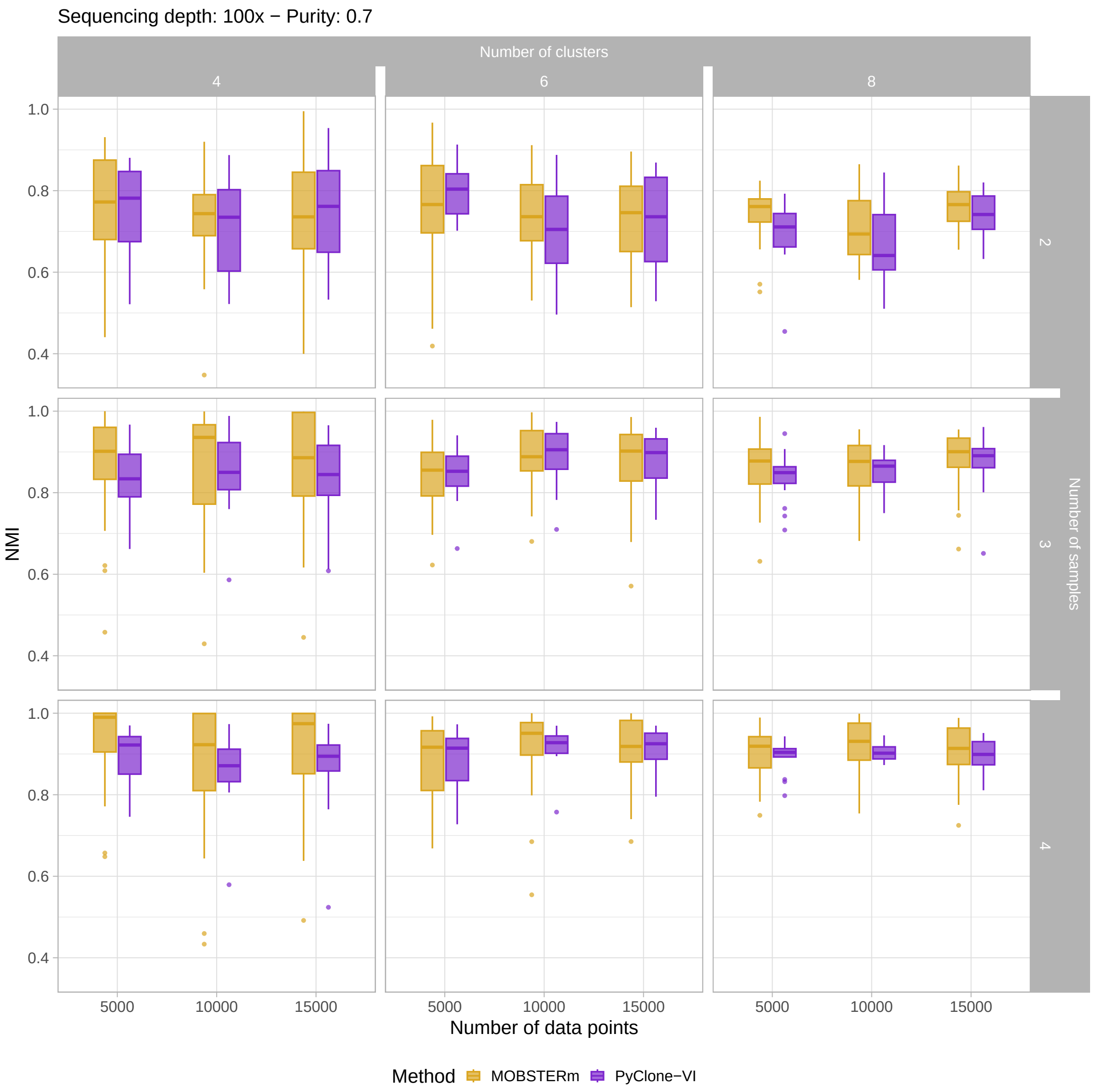

Supplementary Fig. 5: NMI comparison between MOBSTERm (orange) and PyClone-VI [1] (violet) for synthetic datasets with purities of 90% and sequencing depth of 100x.

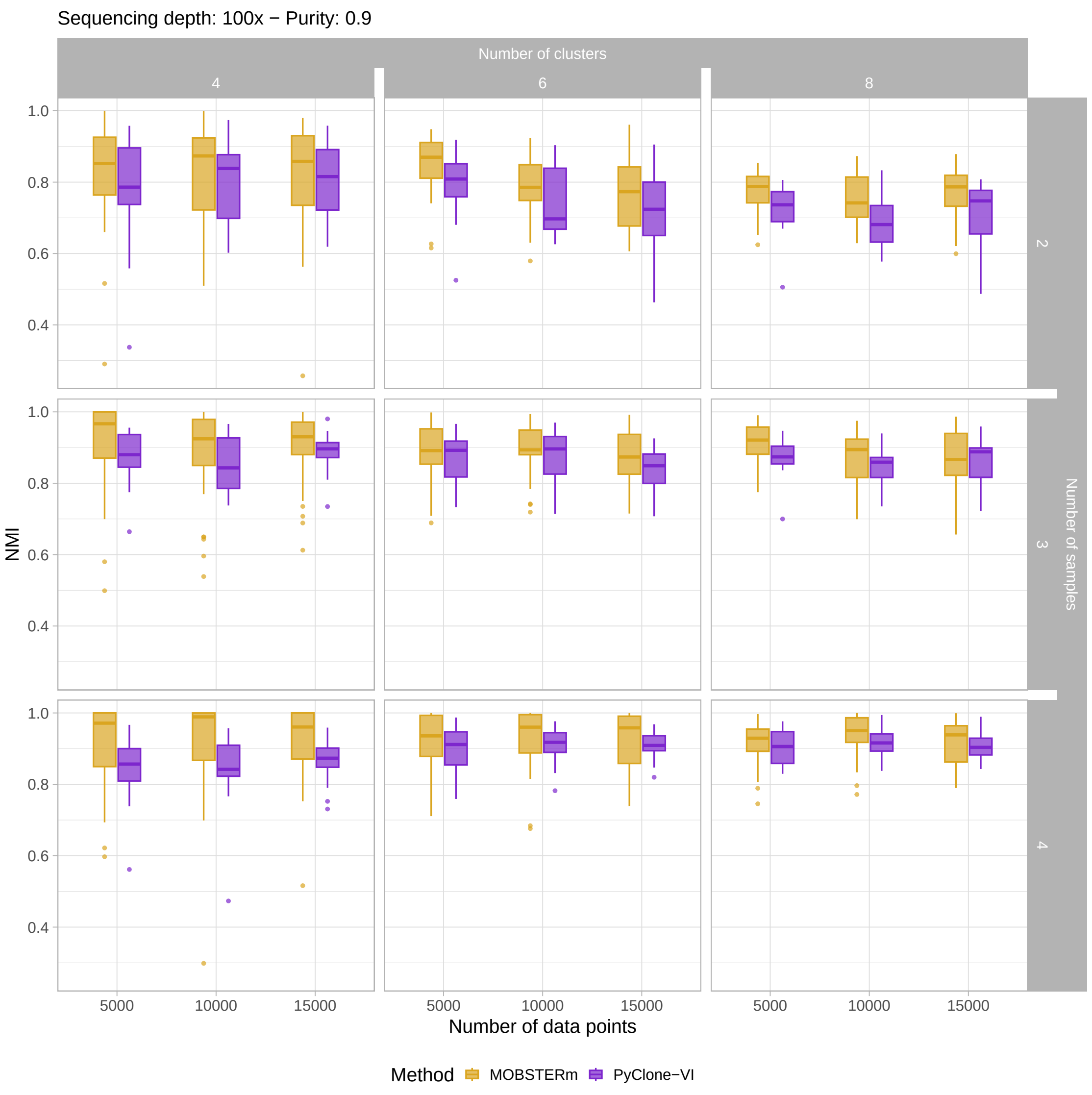

Supplementary Fig. 6: NMI comparison between MOBSTERm (orange) and PyClone-VI [1] (violet) for synthetic datasets with purities of 100% and sequencing depth of 100x.

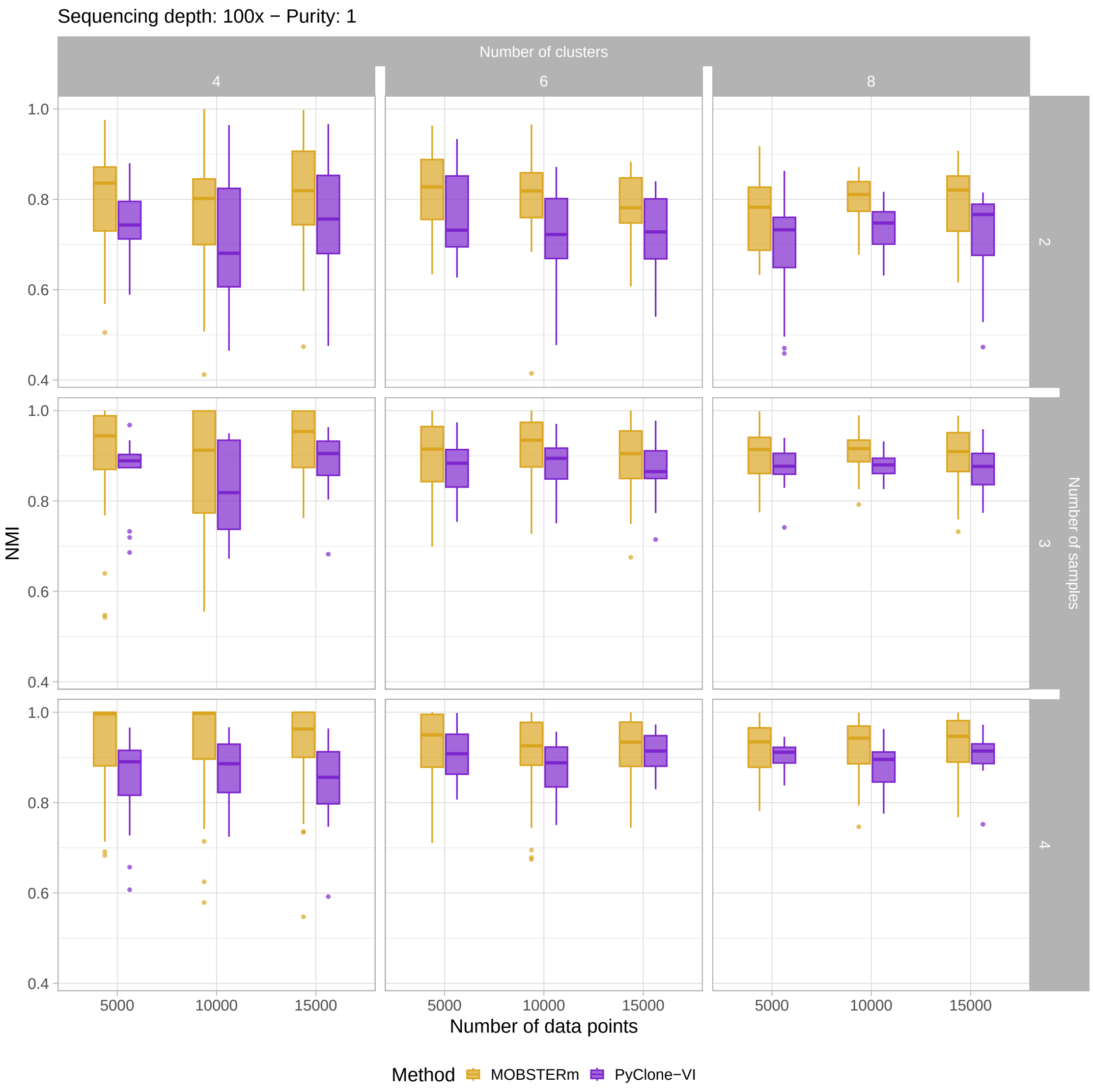

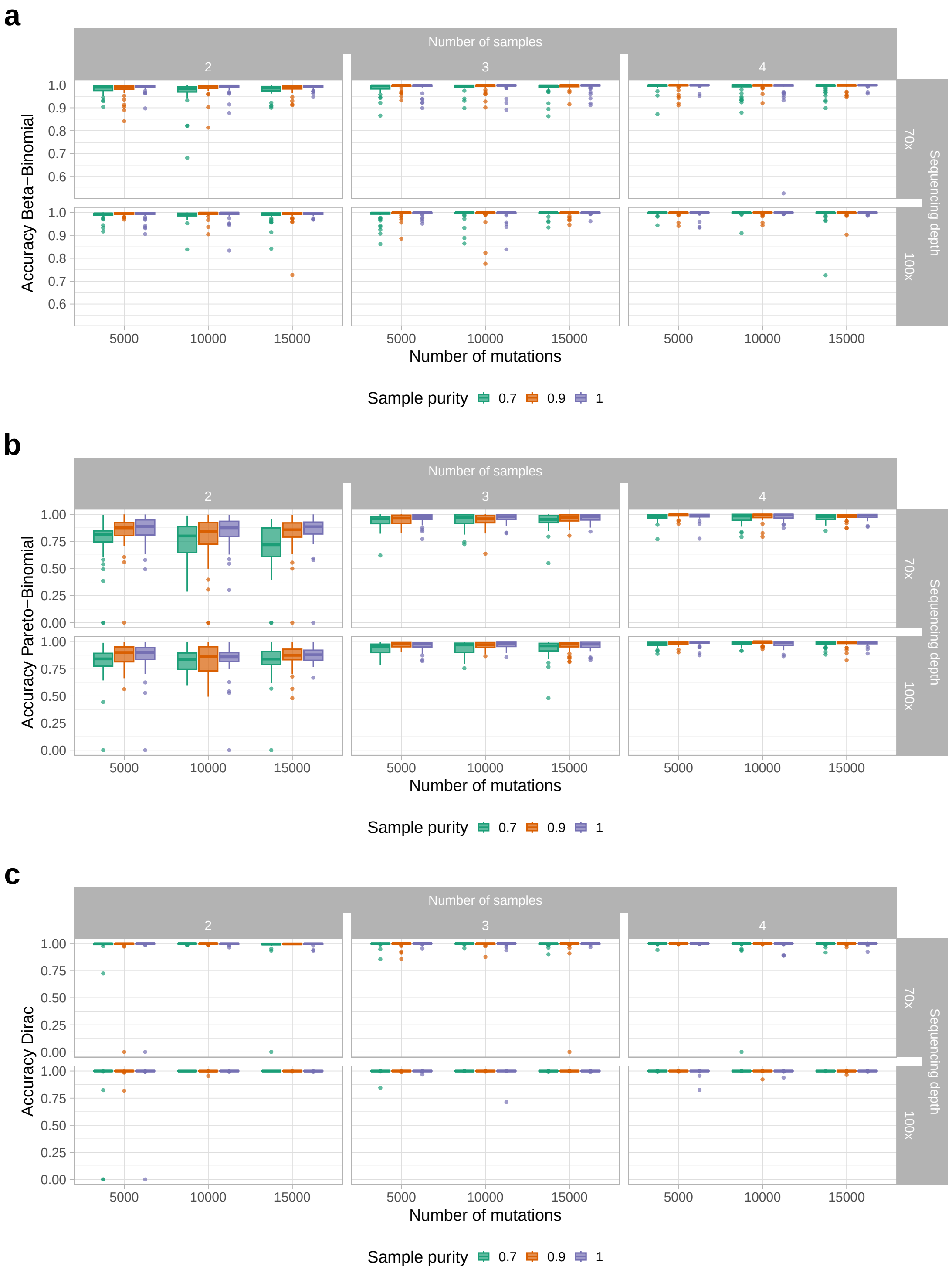


Supplementary Fig. 7: Accuracy of predicted mutation distribution types compared to their true distribution, divided by distribution type (Pareto-Binomial, Beta-Binomial, or Dirac). **a.** Accuracy of predicted distribution types for mutations with a true Beta-Binomial distribution.**b.** Accuracy of predicted distribution types for mutations with a true Pareto-Binomial distribution. **c.** Accuracy of predicted distribution types for mutations with a true Dirac distribution.

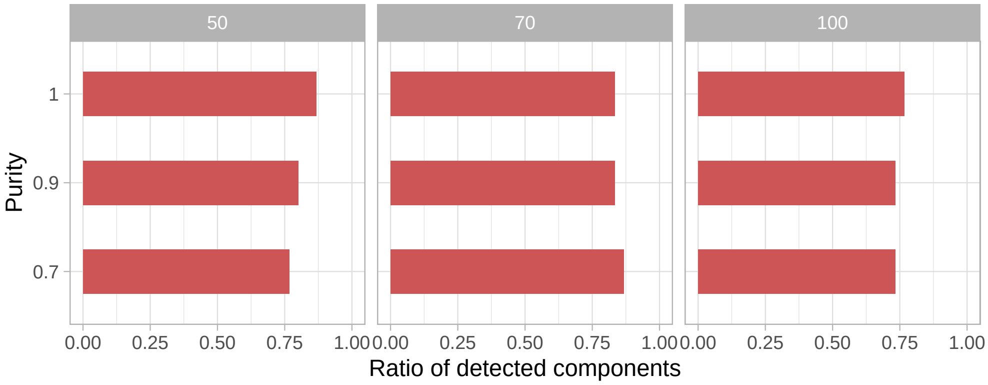


Supplementary Fig. 8: Accuracy in detecting the hitchhiker's mirage in 135 datasets generated with RACES [2], with different purities of 70, 90 and 100% and sequencing depth of 50x, 70x and 100x. The accuracy is computed as the ratio of correctly identified hitchhiker components across all samples.

Supplementary Fig. 9: **Analysis of longitudinal glioblastoma samples**. Multi-page figure for the analysis of 12 glioblastoma patients [3]. **a**. Scatterplot of primary vs relapse VAF. **b**. Primary and relapse VAF. **c**. Heatmap of the $\boldsymbol{\delta}$ parameter, indicating the probabilities assigned to each possible distribution type (Pareto-Binomial, Beta-Binomial, or Dirac) for individual clusters in each sample. **d**. Heatmap of the responsibilities, i.e. posterior assignment probabilities of a data point to a specific cluster given the model parameters. **e**. Mixing proportions. **f**. Number of mutations assigned to each cluster.

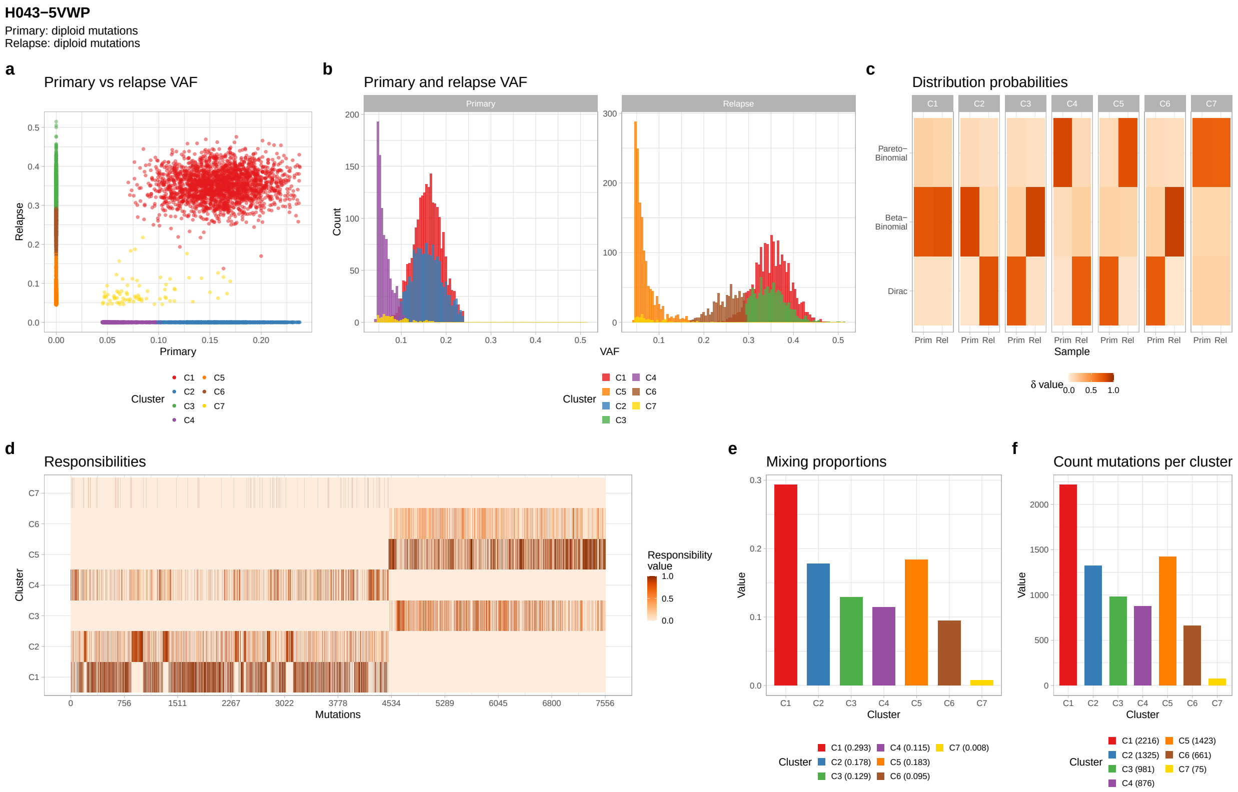

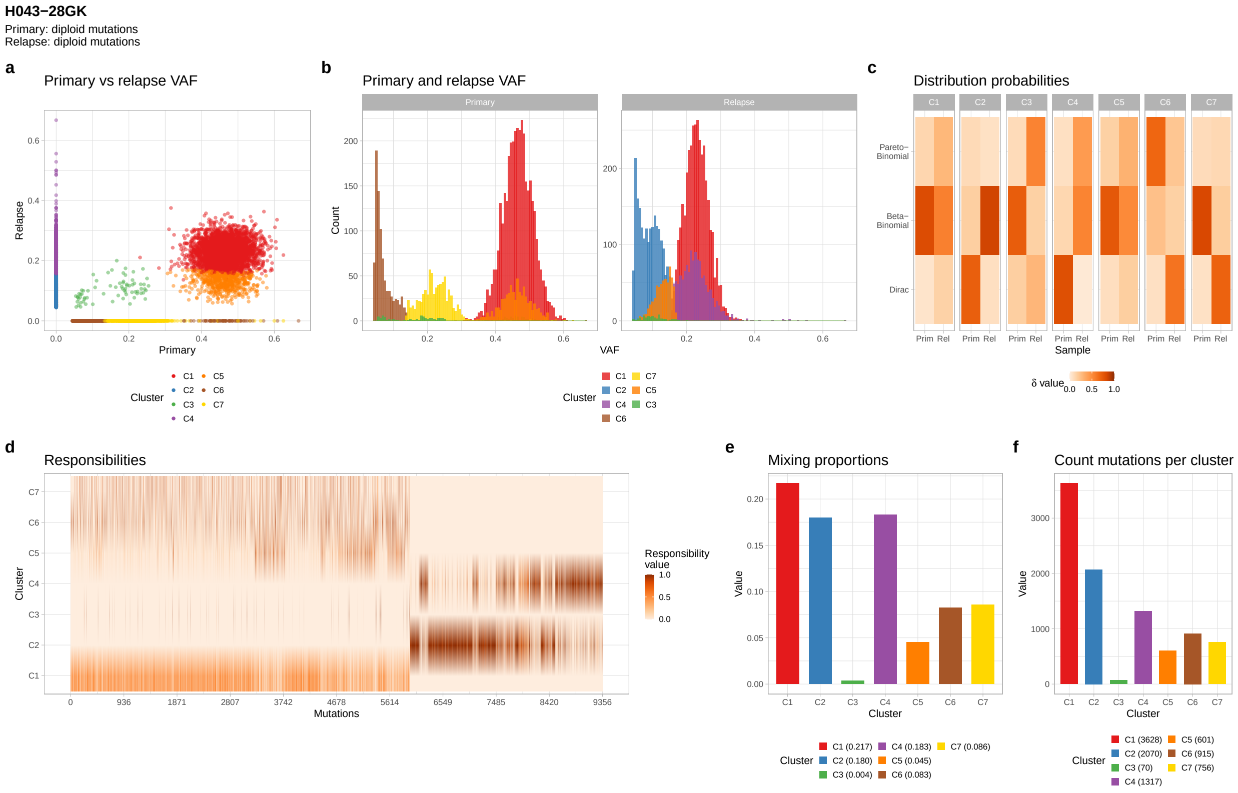
H043−28GK
Primary: diploid mutations
Relapse: diploid mutations
a
Primary vs relapse VAF
Relapse
Primary
Cluster
C1 C2 C3 C4 C5 C6 C7
b
Primary and relapse VAF
Primary
Relapse
Count
VAF
Cluster
C1 C2 C4 C6 C7 C5 C3
c
Distribution probabilities
C1 C2 C3 C4 C5 C6 C7
Pareto−Binomial
Beta−Binomial
Dirac
Prim Rel
Sample
δ value
0.0 0.5 1.0
d
Responsibilities
Cluster
C7 C6 C5 C4 C3 C2 C1
0 936 1871 2807 3742 4678 5614 6549 7485 8420 9356
Mutations
Responsibility value
1.0 0.5 0.0
e
Mixing proportions
Value
Cluster
Cluster
C1 (0.217) C2 (0.180) C3 (0.004) C4 (0.183) C5 (0.045) C6 (0.083) C7 (0.086)
f
Count mutations per cluster
Value
Cluster
Cluster
C1 (3628) C2 (2070) C3 (70) C4 (1317) C5 (601) C6 (915) C7 (756)

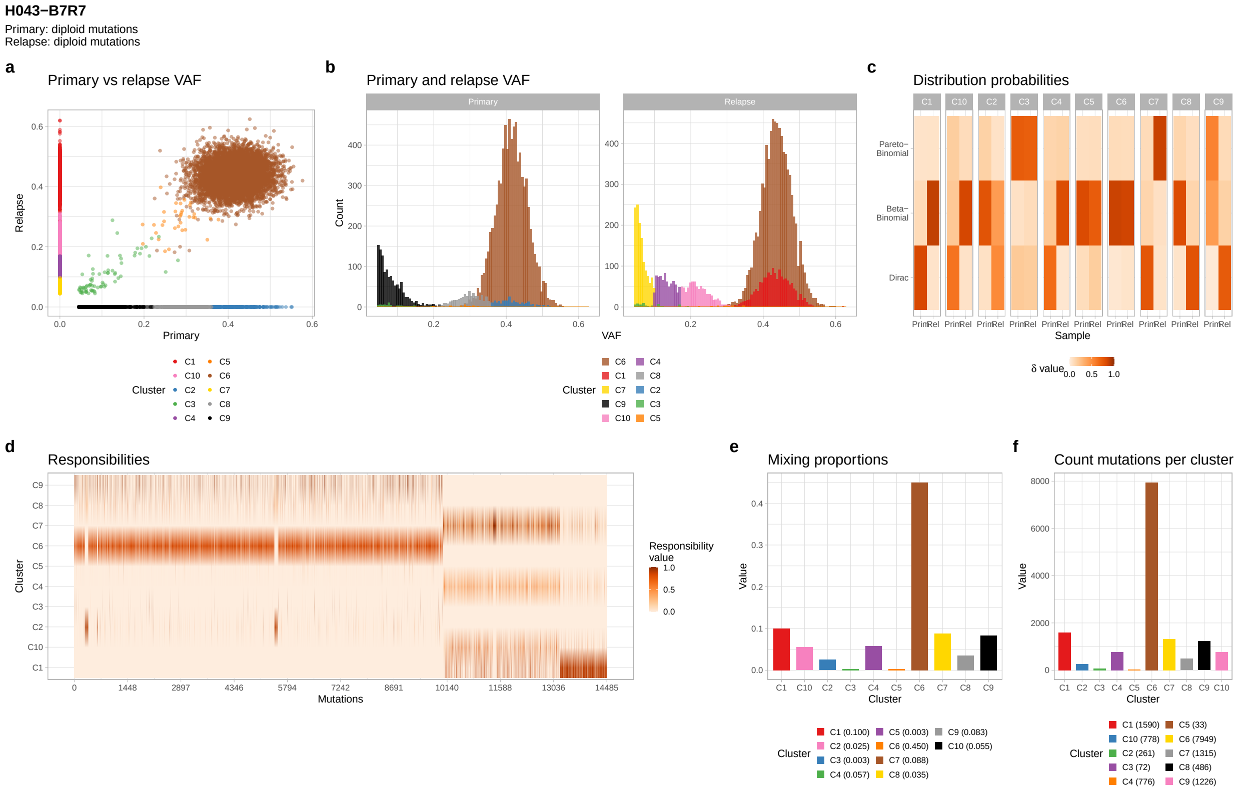
H043−B7R7
Primary: diploid mutations
Relapse: diploid mutations
a
Primary vs relapse VAF
Relapse
Primary
Cluster
C1 C10 C2 C3 C4 C5 C6 C7 C8 C9
b
Primary and relapse VAF
Primary
Relapse
Count
VAF
Cluster
C6 C1 C7 C9 C10 C4 C8 C2 C3 C5
c
Distribution probabilities
C1 C10 C2 C3 C4 C5 C6 C7 C8 C9
Pareto−Binomial
Beta−Binomial
Dirac
Prim Rel
Sample
δ value
0.0 0.5 1.0
d
Responsibilities
Cluster
C9 C8 C7 C6 C5 C4 C3 C2 C10 C1
0 1448 2897 4346 5794 7242 8691 10140 11588 13036 14485
Mutations
Responsibility value
1.0 0.5 0.0
e
Mixing proportions
Value
Cluster
Cluster
C1 (0.100) C2 (0.025) C3 (0.003) C4 (0.057) C5 (0.003) C6 (0.450) C7 (0.088) C8 (0.035) C9 (0.083) C10 (0.055)
f
Count mutations per cluster
Value
Cluster
Cluster
C1 (1590) C10 (778) C2 (261) C3 (72) C4 (776) C5 (33) C6 (7949) C7 (1315) C8 (486) C9 (1226)

**H043−BU96**
Primary: diploid mutations
Relapse: diploid mutations

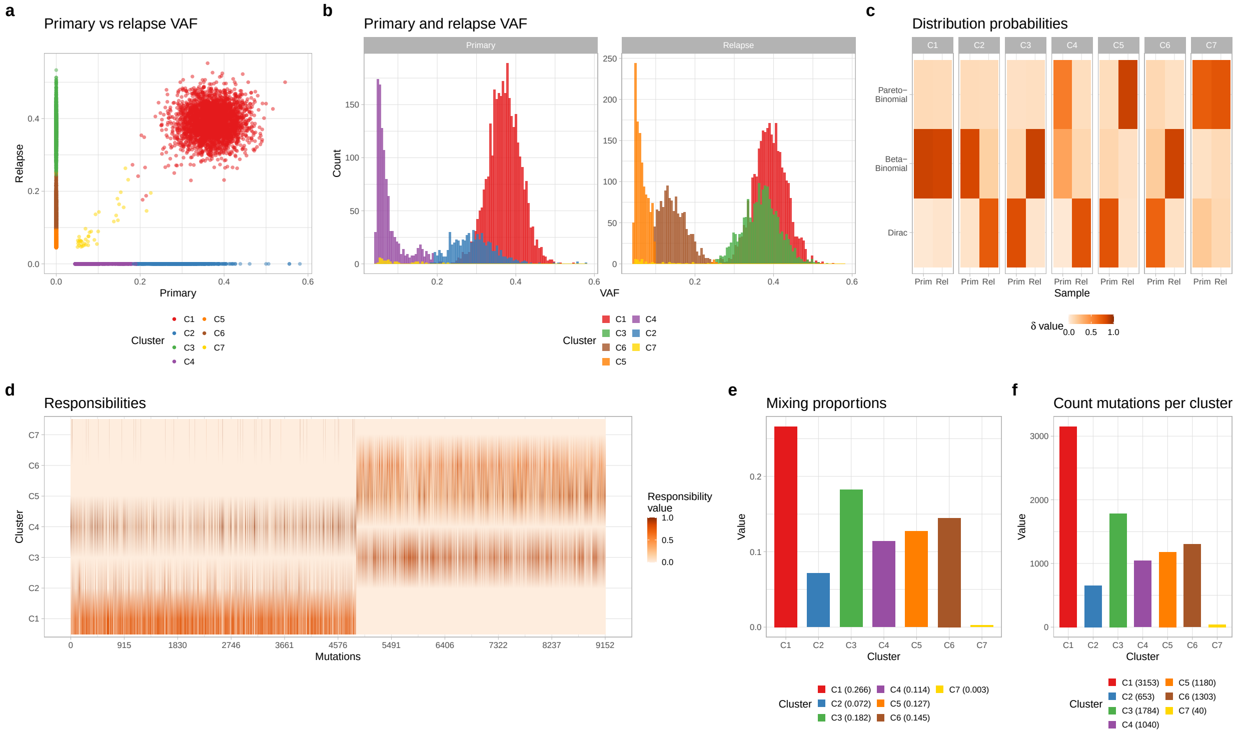


**H043-D9MRCY**
Primary: diploid mutations
Relapse: diploid mutations

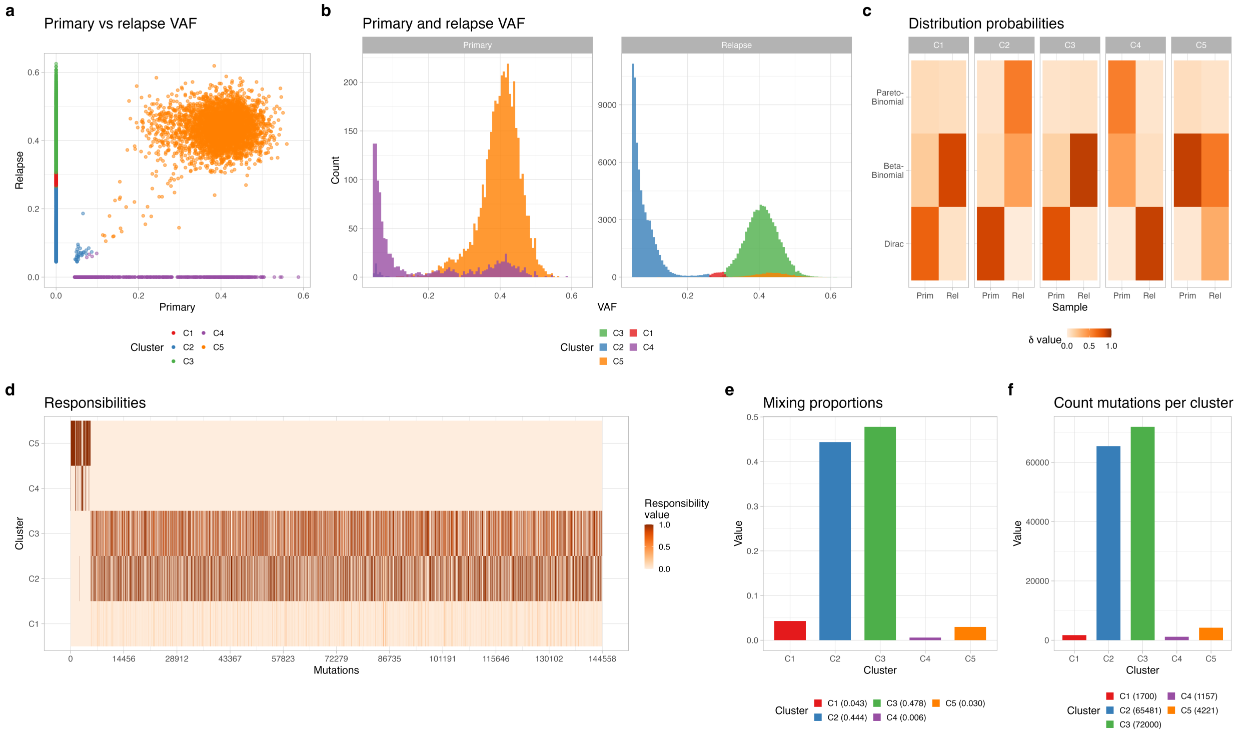

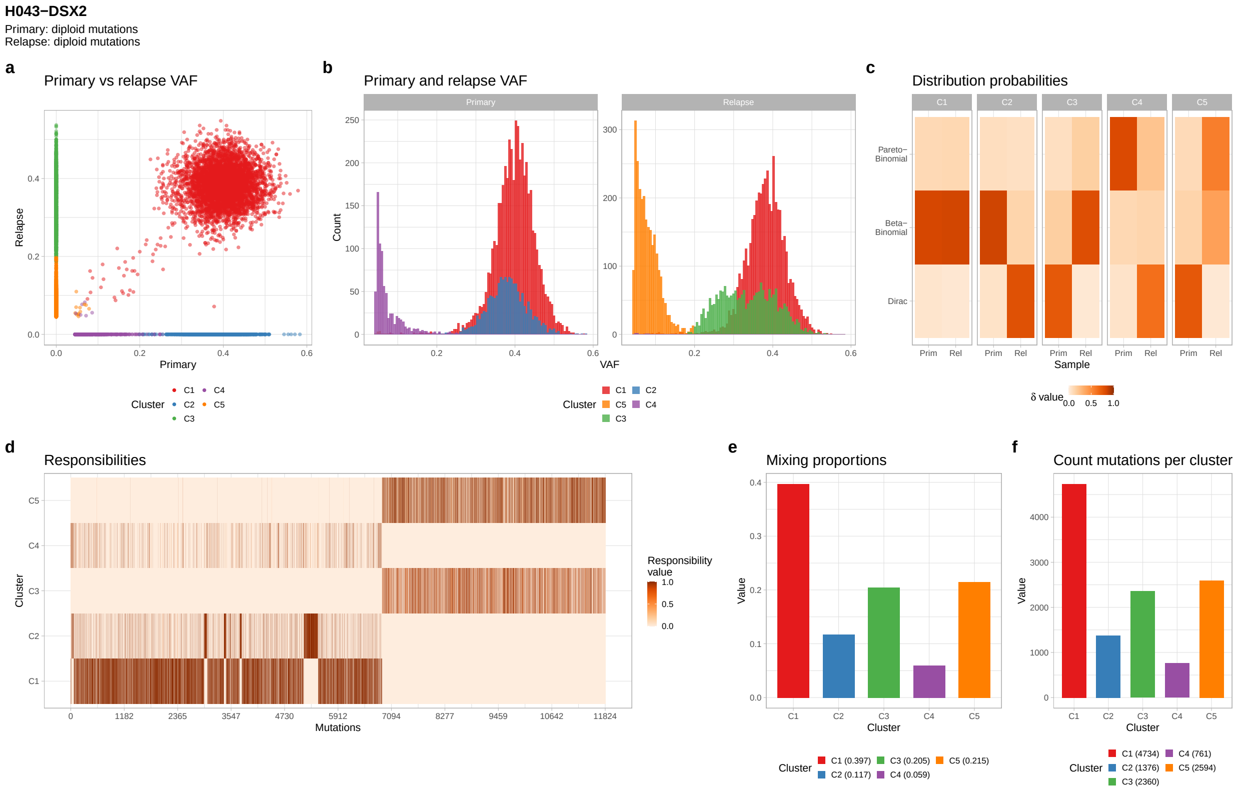
H043−DSX2
Primary: diploid mutations
Relapse: diploid mutations
a
Primary vs relapse VAF
Relapse
Primary
Cluster C1 C2 C3 C4 C5
b
Primary and relapse VAF
Primary
Relapse
Count
VAF
Cluster C1 C5 C3 C2 C4
c
Distribution probabilities
C1 C2 C3 C4 C5
Pareto−Binomial
Beta−Binomial
Dirac
Prim Rel
Sample
δ value 0.0 0.5 1.0
d
Responsibilities
C5
C4
C3
C2
C1
Cluster
Mutations
Responsibility value
1.0
0.5
0.0
e
Mixing proportions
Value
Cluster
Cluster C1 (0.397) C2 (0.117) C3 (0.205) C4 (0.059) C5 (0.215)
f
Count mutations per cluster
Value
Cluster
Cluster C1 (4734) C2 (1376) C3 (2360) C4 (761) C5 (2594)

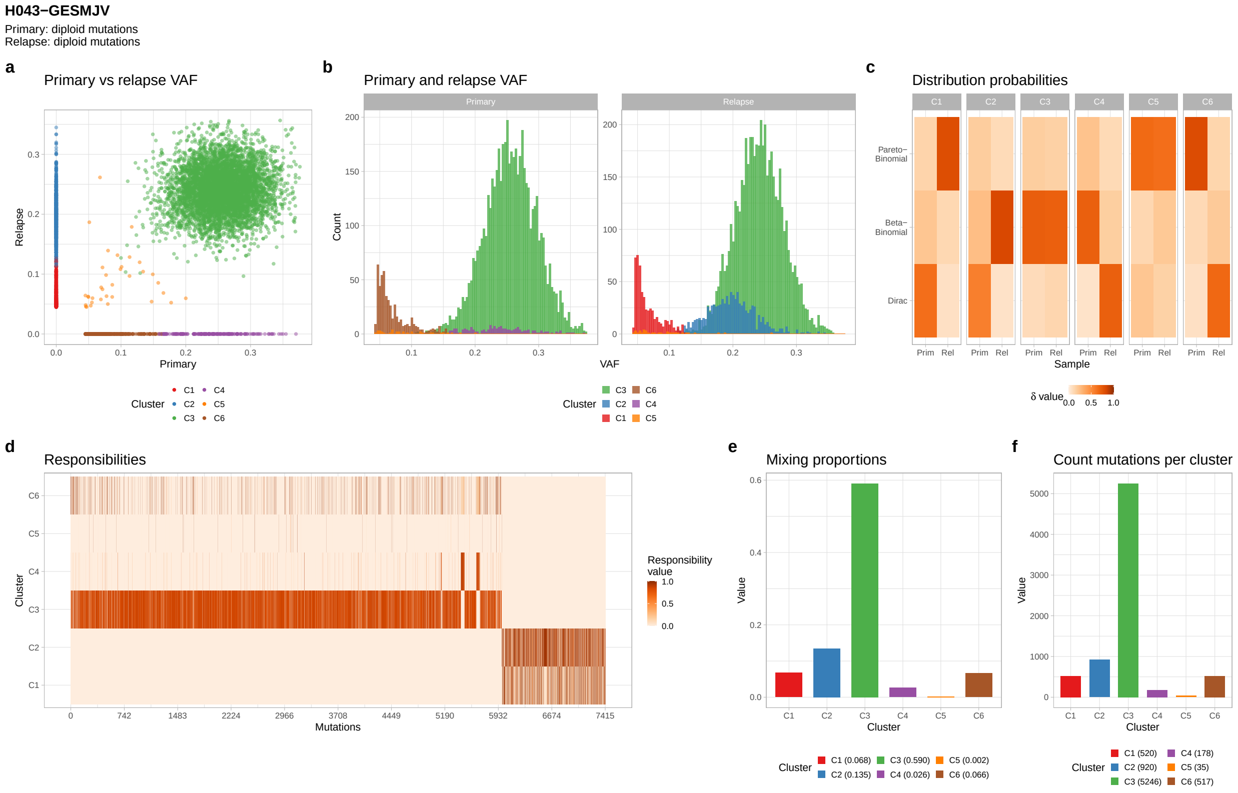
H043−GESMJV
Primary: diploid mutations
Relapse: diploid mutations
a
Primary vs relapse VAF
Relapse
Primary
Cluster C1 C2 C3 C4 C5 C6
b
Primary and relapse VAF
Primary
Relapse
Count
VAF
Cluster C3 C2 C1 C6 C4 C5
c
Distribution probabilities
C1 C2 C3 C4 C5 C6
Pareto−Binomial
Beta−Binomial
Dirac
Prim Rel
Sample
δ value 0.0 0.5 1.0
d
Responsibilities
C6
C5
C4
C3
C2
C1
Cluster
Mutations
Responsibility value
1.0
0.5
0.0
e
Mixing proportions
Value
Cluster
Cluster C1 (0.068) C2 (0.135) C3 (0.590) C4 (0.026) C5 (0.002) C6 (0.066)
f
Count mutations per cluster
Value
Cluster
Cluster C1 (520) C2 (920) C3 (5246) C4 (178) C5 (35) C6 (517)

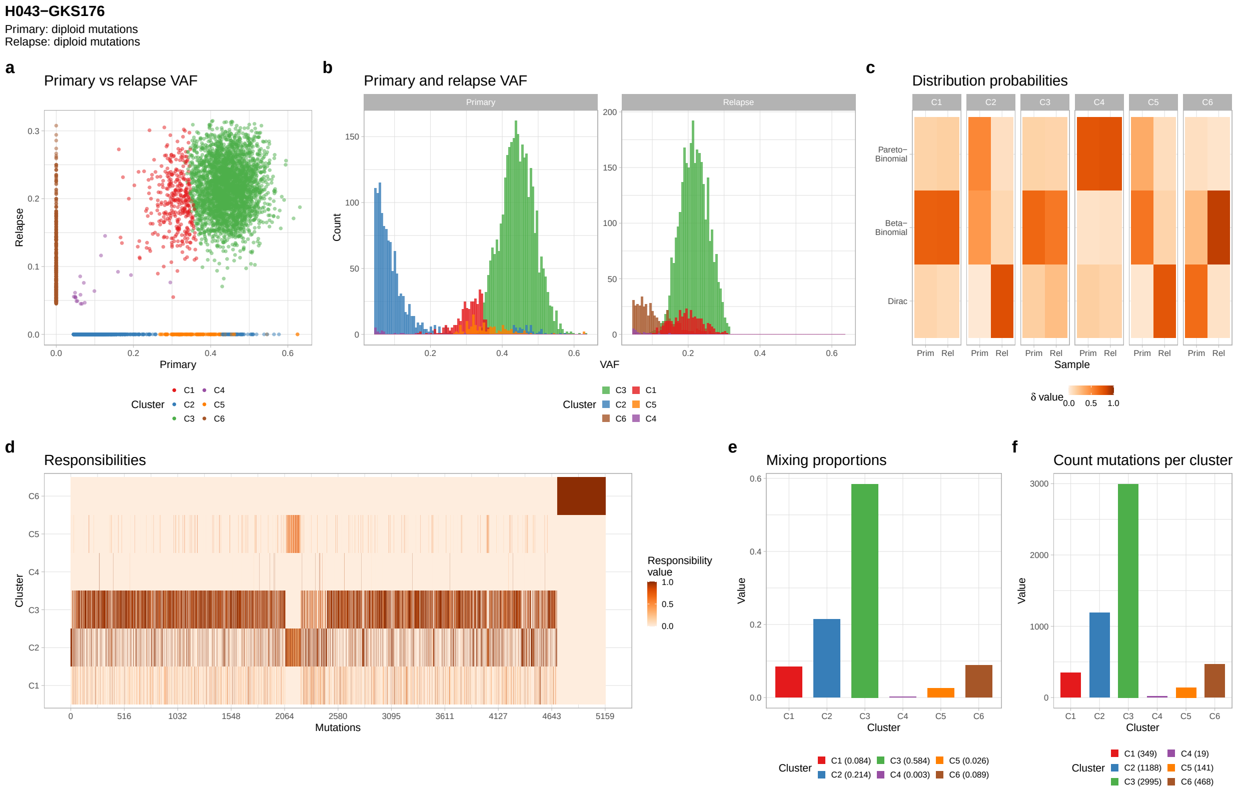
H043−GKS176
Primary: diploid mutations
Relapse: diploid mutations
a
Primary vs relapse VAF
Relapse
Primary
Cluster C1 C2 C3 C4 C5 C6
b
Primary and relapse VAF
Primary
Relapse
Count
VAF
Cluster C3 C2 C6 C1 C5 C4
c
Distribution probabilities
C1 C2 C3 C4 C5 C6
Pareto−Binomial
Beta−Binomial
Dirac
Prim Rel
Sample
δ value 0.0 0.5 1.0
d
Responsibilities
Cluster
Mutations
Responsibility value 1.0 0.5 0.0
e
Mixing proportions
Value
Cluster
C1 (0.084) C2 (0.214) C3 (0.584) C4 (0.003) C5 (0.026) C6 (0.089)
f
Count mutations per cluster
Value
Cluster
C1 (349) C2 (1188) C3 (2995) C4 (19) C5 (141) C6 (468)

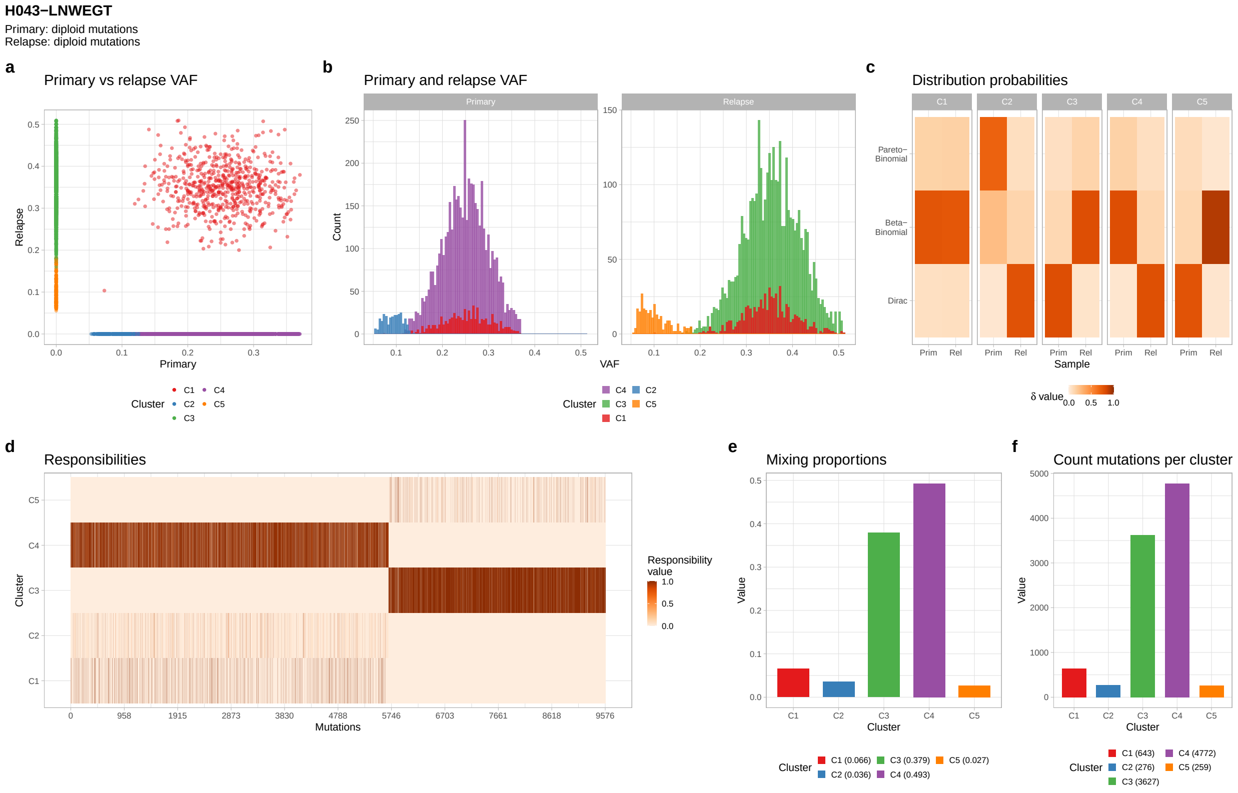
H043−LNWEGT
Primary: diploid mutations
Relapse: diploid mutations
a
Primary vs relapse VAF
Relapse
Primary
Cluster C1 C2 C3 C4 C5
b
Primary and relapse VAF
Primary
Relapse
Count
VAF
Cluster C4 C3 C1 C2 C5
c
Distribution probabilities
C1 C2 C3 C4 C5
Pareto−Binomial
Beta−Binomial
Dirac
Prim Rel
Sample
δ value 0.0 0.5 1.0
d
Responsibilities
Cluster
Mutations
Responsibility value 1.0 0.5 0.0
e
Mixing proportions
Value
Cluster
C1 (0.066) C2 (0.036) C3 (0.379) C4 (0.493) C5 (0.027)
f
Count mutations per cluster
Value
Cluster
C1 (643) C2 (276) C3 (3627) C4 (4772) C5 (259)

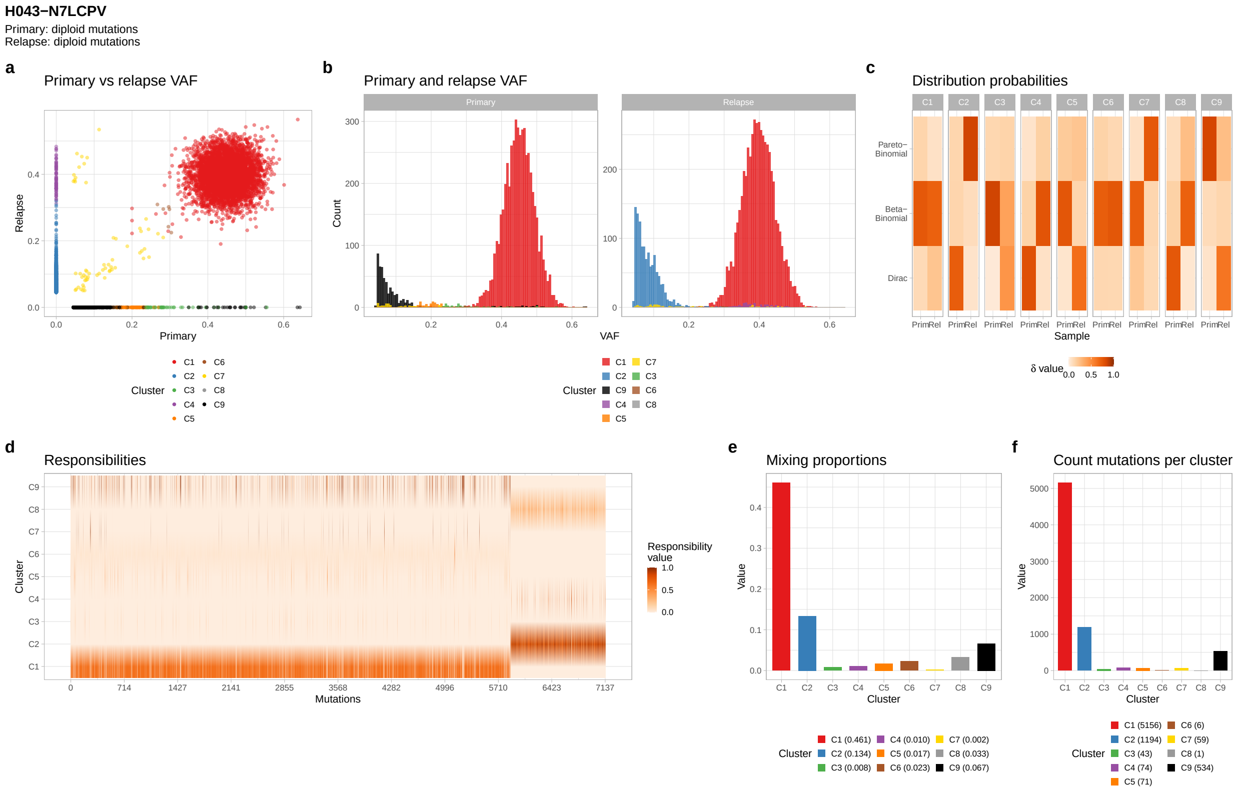

H043−N7LCPV
Primary: diploid mutations
Relapse: diploid mutations
a
Primary vs relapse VAF
Relapse
Primary
Cluster
C1 C2 C3 C4 C5 C6 C7 C8 C9
b
Primary and relapse VAF
Primary
Relapse
Count
VAF
Cluster
C1 C2 C9 C4 C5 C7 C3 C6 C8
c
Distribution probabilities
Pareto−Binomial
Beta−Binomial
Dirac
Prim Rel
Sample
δ value
d
Responsibilities
Cluster
Mutations
Responsibility value
e
Mixing proportions
Value
Cluster
C1 (0.461) C2 (0.134) C3 (0.008) C4 (0.010) C5 (0.017) C6 (0.023) C7 (0.002) C8 (0.033) C9 (0.067)
f
Count mutations per cluster
Value
Cluster
C1 (5156) C2 (1194) C3 (43) C4 (74) C5 (71) C6 (6) C7 (59) C8 (1) C9 (534)


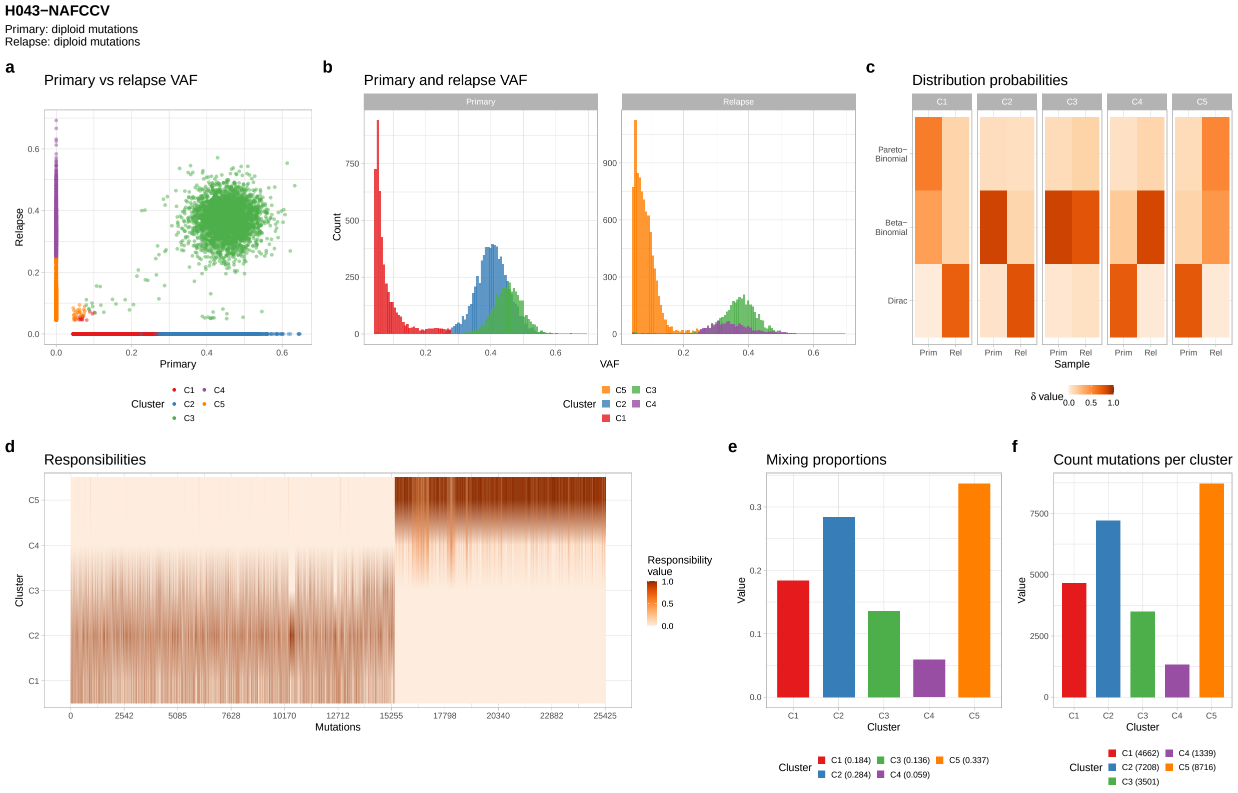

H043−NAFCCV
Primary: diploid mutations
Relapse: diploid mutations
a
Primary vs relapse VAF
Relapse
Primary
Cluster
C1 C2 C3 C4 C5
b
Primary and relapse VAF
Primary
Relapse
Count
VAF
Cluster
C5 C2 C1 C3 C4
c
Distribution probabilities
Pareto−Binomial
Beta−Binomial
Dirac
Prim Rel
Sample
δ value
d
Responsibilities
Cluster
Mutations
Responsibility value
e
Mixing proportions
Value
Cluster
C1 (0.184) C2 (0.284) C3 (0.136) C4 (0.059) C5 (0.337)
f
Count mutations per cluster
Value
Cluster
C1 (4662) C2 (7208) C3 (3501) C4 (1339) C5 (8716)

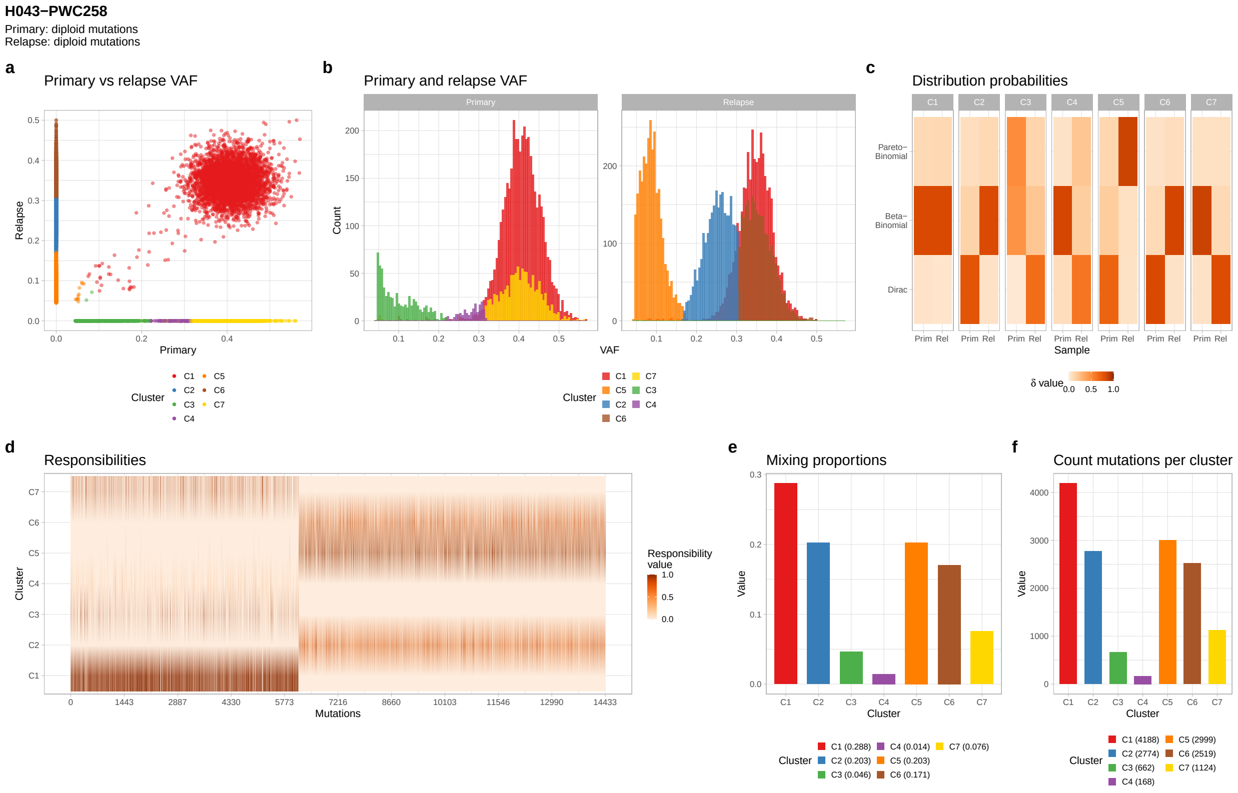
H043−PWC258
Primary: diploid mutations
Relapse: diploid mutations
a
Primary vs relapse VAF
Relapse
Primary
Cluster
C1 C2 C3 C4 C5 C6 C7
b
Primary and relapse VAF
Primary
Relapse
Count
VAF
Cluster
C1 C5 C2 C6 C7 C3 C4
c
Distribution probabilities
C1 C2 C3 C4 C5 C6 C7
Pareto−Binomial
Beta−Binomial
Dirac
Prim Rel
Sample
δ value
0.0 0.5 1.0
d
Responsibilities
Cluster
Mutations
0 1443 2887 4330 5773 7216 8660 10103 11546 12990 14433
Responsibility value
1.0 0.5 0.0
e
Mixing proportions
Value
Cluster
C1 (0.288) C2 (0.203) C3 (0.046) C4 (0.014) C5 (0.203) C6 (0.171) C7 (0.076)
f
Count mutations per cluster
Value
Cluster
C1 (4188) C2 (2774) C3 (662) C4 (168) C5 (2999) C6 (2519) C7 (1124)

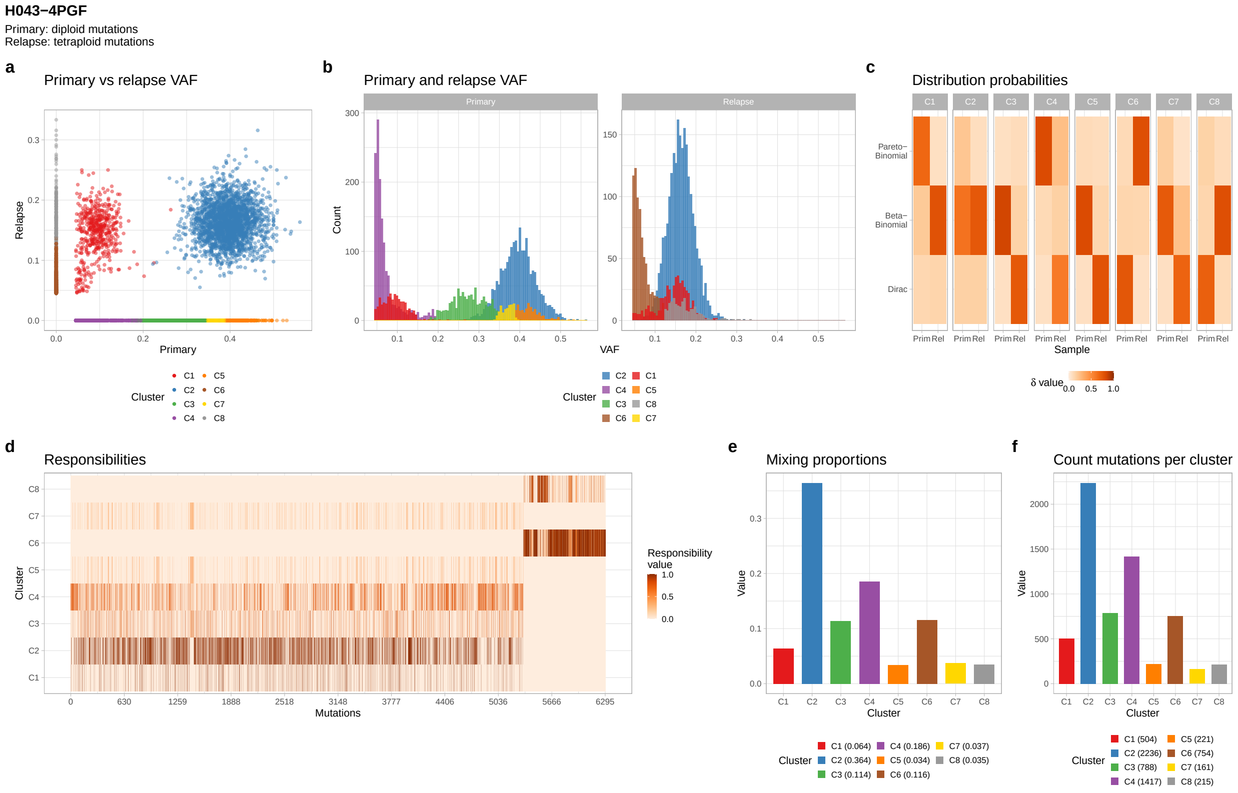
H043−4PGF
Primary: diploid mutations
Relapse: tetraploid mutations
a
Primary vs relapse VAF
Relapse
Primary
Cluster
C1 C2 C3 C4 C5 C6 C7 C8
b
Primary and relapse VAF
Primary
Relapse
Count
VAF
Cluster
C2 C4 C3 C6 C1 C5 C8 C7
c
Distribution probabilities
C1 C2 C3 C4 C5 C6 C7 C8
Pareto−Binomial
Beta−Binomial
Dirac
Prim Rel
Sample
δ value
0.0 0.5 1.0
d
Responsibilities
Cluster
Mutations
0 630 1259 1888 2518 3148 3777 4406 5036 5666 6295
Responsibility value
1.0 0.5 0.0
e
Mixing proportions
Value
Cluster
C1 (0.064) C2 (0.364) C3 (0.114) C4 (0.186) C5 (0.034) C6 (0.116) C7 (0.037) C8 (0.035)
f
Count mutations per cluster
Value
Cluster
C1 (504) C2 (2236) C3 (788) C4 (1417) C5 (221) C6 (754) C7 (161) C8 (215)

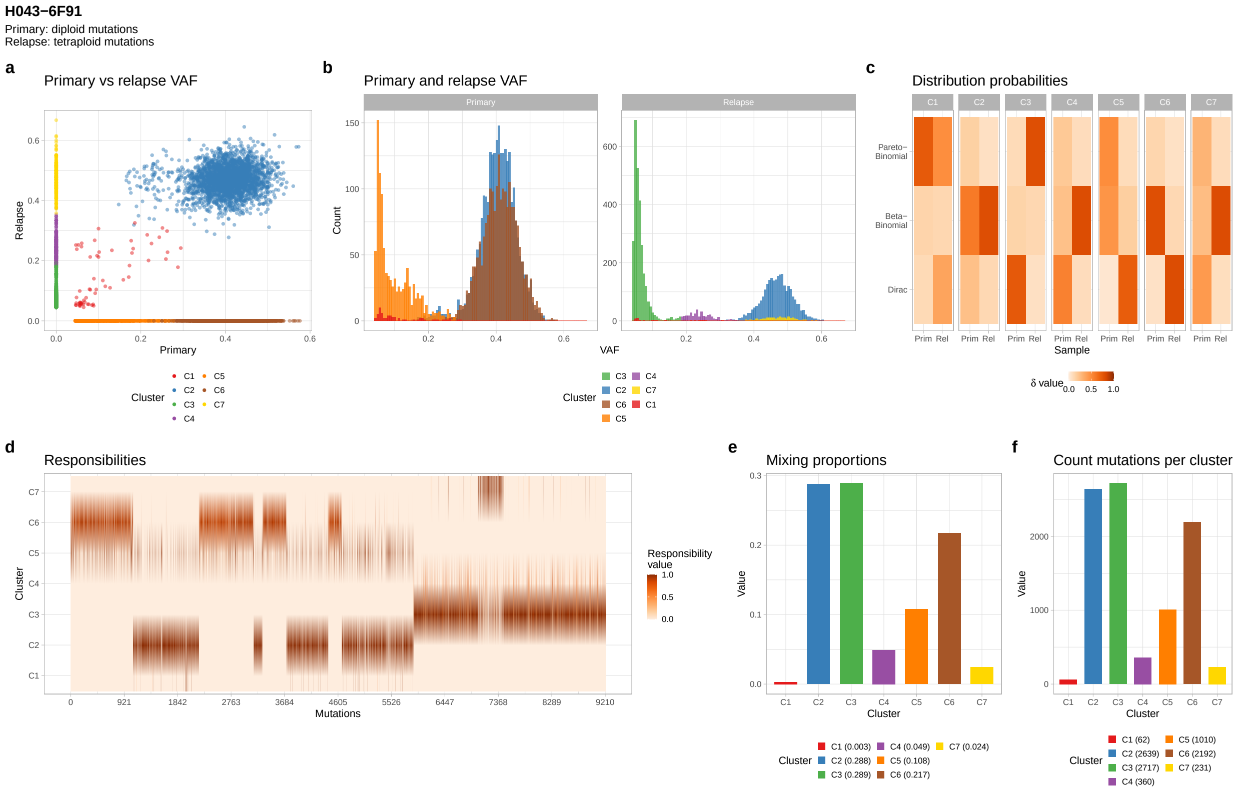
H043−6F91
Primary: diploid mutations
Relapse: tetraploid mutations
a
Primary vs relapse VAF
Relapse
Primary
Cluster
C1 C2 C3 C4 C5 C6 C7
b
Primary and relapse VAF
Primary
Relapse
Count
VAF
Cluster
C3 C2 C6 C5 C4 C7 C1
c
Distribution probabilities
C1 C2 C3 C4 C5 C6 C7
Pareto−Binomial
Beta−Binomial
Dirac
Prim Rel
Sample
δ value
0.0 0.5 1.0
d
Responsibilities
Cluster
Mutations
Responsibility value
1.0 0.5 0.0
e
Mixing proportions
Value
Cluster
C1 (0.003) C2 (0.288) C3 (0.289) C4 (0.049) C5 (0.108) C6 (0.217) C7 (0.024)
f
Count mutations per cluster
Value
Cluster
C1 (62) C2 (2639) C3 (2717) C4 (360) C5 (1010) C6 (2192) C7 (231)

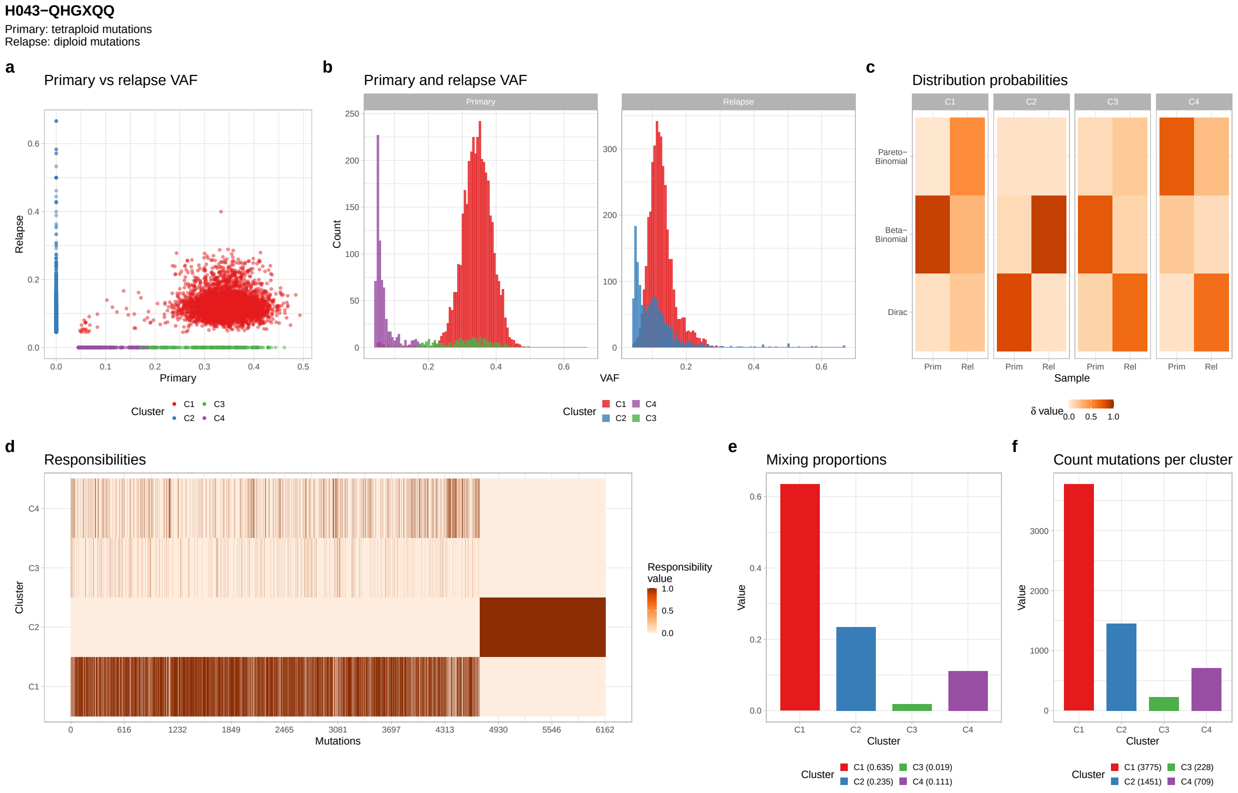
H043−QHGXQQ
Primary: tetraploid mutations
Relapse: diploid mutations
a
Primary vs relapse VAF
Relapse
Primary
Cluster
C1 C2 C3 C4
b
Primary and relapse VAF
Primary
Relapse
Count
VAF
Cluster
C1 C2 C4 C3
c
Distribution probabilities
C1 C2 C3 C4
Pareto−Binomial
Beta−Binomial
Dirac
Prim Rel
Sample
δ value
0.0 0.5 1.0
d
Responsibilities
Cluster
Mutations
Responsibility value
1.0 0.5 0.0
e
Mixing proportions
Value
Cluster
C1 (0.635) C2 (0.235) C3 (0.019) C4 (0.111)
f
Count mutations per cluster
Value
Cluster
C1 (3775) C2 (1451) C3 (228) C4 (709)

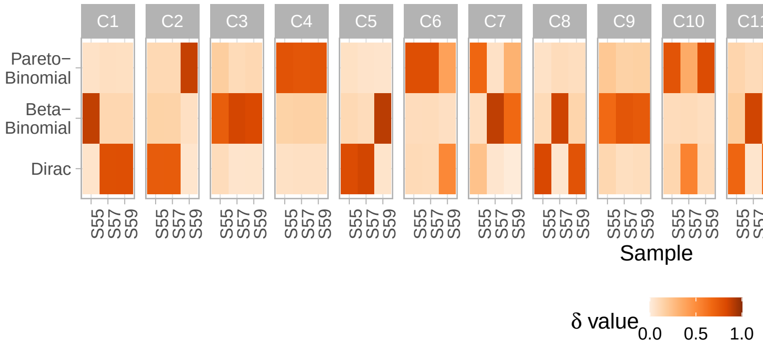


Supplementary Fig. 10: Heatmap of the $\boldsymbol{\delta}$ parameter for samples S55, S57 and S59 for all clusters of Set07.
